# Bridging Glucose Metabolism and Intrinsic Functional Organization of the Human Cortex

**DOI:** 10.1101/2024.09.26.615152

**Authors:** Bin Wan, Valentin Riedl, Gabriel Castrillon, Matthias Kirschner, Sofie L. Valk

## Abstract

The human brain requires a continuous supply of energy to function effectively. Here, we investigated how the low-dimensional organization of intrinsic functional connectivity patterns based on resting-state functional magnetic resonance imaging relates to brain energy expenditure measured by fluorodeoxyglucose positron emission tomography. By incrementally adding more dimensions of brain organization (via functional gradients), we were able to show that increasing amounts of variance in the map of brain energy expenditure could be accounted for. In particular, the brain organization dimensions that explained a large amount of the variance in intrinsic brain function also explained a large amount of the regional variance in the energy expenditure maps. This was particularly true for brain organization maps based on the strongest connections, suggesting that "weak" connections may not explain as much energy variance. Notably, our topological model was more effective than random brain organization configurations, suggesting that brain organization may be specifically associated with energy optimization. Finally, using brain asymmetry as a model for metabolic efficiency, we found that optimizing energy expenditure independently in each hemisphere outperformed non-independent optimization. This supports the concept of hemispheric competition rather than lateralization in energy allocation. Our results demonstrate how the spatial organization of functional connections is systematically linked to optimized energy expenditure in the human brain, providing new insights into the metabolic basis of brain function.

## Main

The human brain has rich patterns of neural activity that consume a considerable amount of metabolic energy and computational resources at rest (Howarth et al., 2012; Hyder et al., 2013; Lennie, 2003; Niven et al., 2007; Shen et al., 1999; Sibson et al., 1998). Previous studies have correlated cortical energy cost with regional features of resting state functional connectome and found that regions with higher local connectivity, centrality, and amplitude of low frequency fluctuations consume more energy (Aiello et al., 2015; Castrillon et al., 2023; Palombit et al., 2022; Riedl et al., 2014; Stiernman et al., 2021; Wang et al., 2021). Although these indices provide local or network graph information about connectivity patterns, these models only partially explain the spatial pattern of energy expenditure. Another way to understand brain function is to conceptualize it as an outcome of inter-regional connectivity, systematically organized by network community integration and segregation, which can be captured by gradients (Margulies et al., 2016). We hypothesize that energy consumption relates to the topological organization of intrinsic brain function.

Positron emission tomography (PET) and functional magnetic resonance imaging (fMRI) are suitable tools for capturing cortical energy expenditure and spatial functional organization (Magistretti et al., 1999; Magistretti & Allaman, 2015; Raichle & Gusnard, 2002). In particular, the glucose analog F18 labeled deoxyglucose (18F-FDG) is a marker of the intracellular glycolytic rate. Due to a lack of a 2-OH group, it cannot be further metabolized along the glycolytic pathway. PET imaging can capture the spatial distribution of accumulated F18 which then allows measurement of regional differences in the cerebral metabolic rate of glucose uptake *in vivo* (CMRglc) (Castrillon et al., 2023; Wu et al., 2003). FDG-PET is reliable for capturing robust energy expenditure of brain activity as the metabolic activity of brain energy expenditure is constant over time and found reliable using test-retest data (Baumgartner et al., 2018; Goutal et al., 2020; Raichle & Gusnard, 2002). Resting-state fMRI captures the time-varied blood-oxygen-level-dependent (BOLD) signals at rest (Kwong et al., 1992; Ogawa et al., 1990) with regions showing pairwise temporal correlation having strong connectivity, i.e., functional connectivity (FC) (Friston, 1994). The resting state signal is thought to be a combination of metabolic, anatomical, and cognitive signals (Pezzulo et al., 2021). Regions with similar FC profiles form patterns of functional organization that reflect axes of topological organization of interregional integration and segregation, i.e., gradients (Bernhardt et al., 2022; Huntenburg et al., 2018; Margulies et al., 2016; Vos de Wael et al., 2020). The first two gradients in healthy young adults describe axes that differentiate sensory from association regions and somatomotor from visual regions, which overlap with models of information processing hierarchy and microstructural variation (Huntenburg et al., 2018; Margulies et al., 2016; Mesulam, 1998). Functional gradients are reproducible and reliable across subjects/sessions (Hong et al., 2020; Knodt et al., 2023; Vos de Wael et al., 2020).

We hypothesize that there is a relationship between topological organization of intrinsic functional connections and energy expenditure, as 1) the organization of the cortex is likely a result of evolutionary pressures on minimizing energy demands (Bullmore & Sporns, 2012; Castrillon et al., 2023; Kuzawa et al., 2014) and 2) regions that have a similar functional organization likely have a similar energy expenditure (Margulies et al., 2016). However, with FDG-PET, we can only obtain a map of overall energy expenditure within a region, not how much energy functional organization consumes. To overcome this challenge, we model energy expenditure using gradients of functional organization as independent variables. Through evaluating the combined variance explained by each gradient, we aimed to reconstruct the energy expenditure map. This approach allows us to describe how different gradients explain the energy regionally consumed, as measured using FDG-PET. **Figure 1** illustrates the principles of BOLD FC similarity concerning energy and oxygen expenditure. The first aim of our study is to evaluate to what extent functional organization and energy expenditure have shared organizational features. We hypothesize that when using gradient maps of functional organization to model the spatial distribution of the CMRglc map, the model performance should increase step by step with an increasingly detailed description of functional brain organization (e.g. increased number of gradients and variance explained). Though generally only the first three gradients are evaluated and functionally interpreted (Huntenburg et al., 2018; Paquola et al., 2022), we aim for a more exhaustive model and test the first 100 gradients. This choice is motivated by previous work that showed prediction accuracy of behavior improved by including 60 gradients or more (Kong et al., 2023). The variance explained by these gradients may capture brain features with behavioral and physiological relevance (Kong et al., 2023). Here the authors showed that combining these gradients could rival the predictive power of models based on parcellations, ultimately bridging global and regional features of brain organization. Moreover, it will allow us to understand whether energy patterning explained by the gradient model gradually increases with the number of gradients, or reaches ceiling after a few dominant organizational principles.

**Figure 1.**
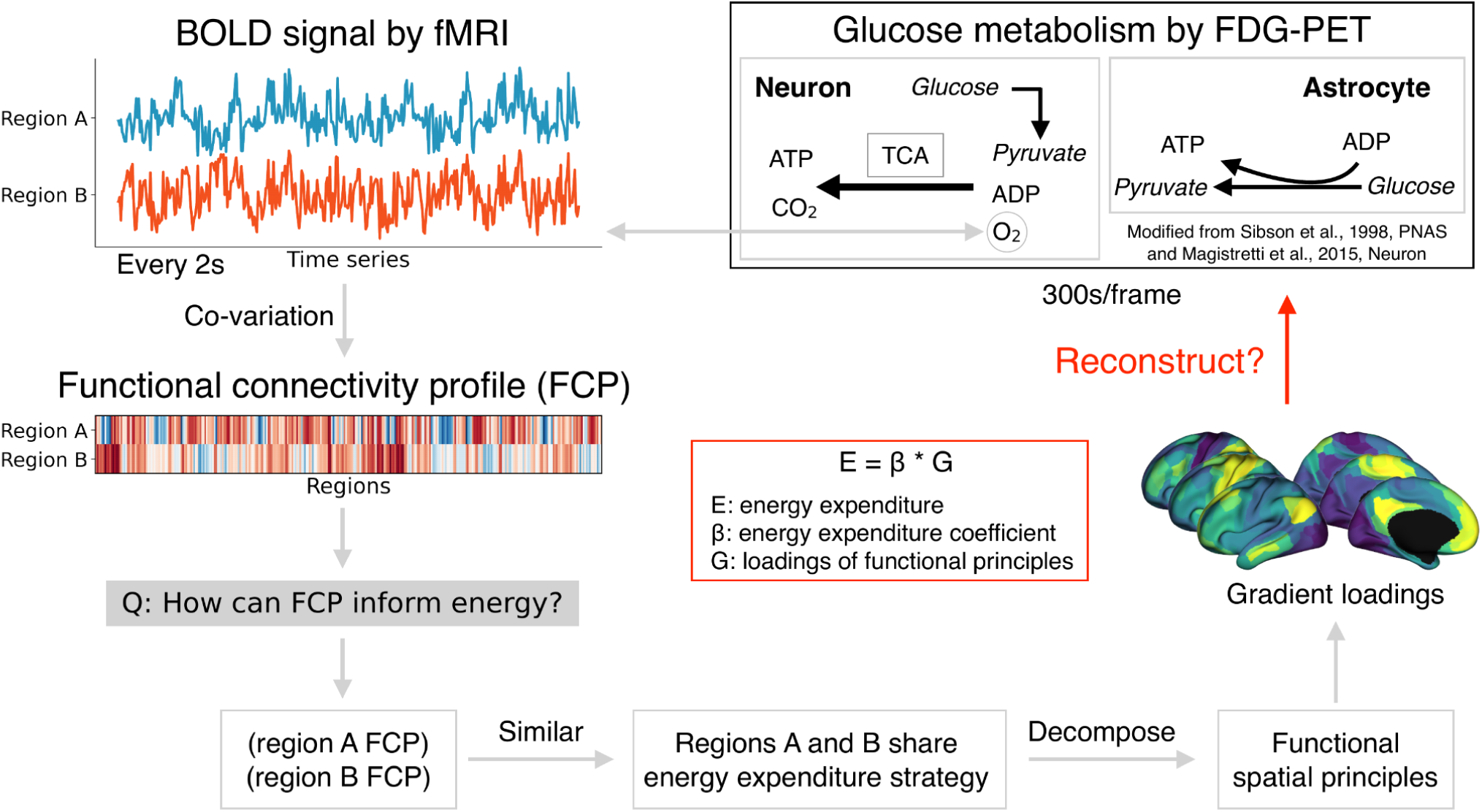
The proposed relationship between glucose metabolism and functional organization.

Importantly, both functional organization and glucose metabolism show hemispheric differences (Berardi et al., 1991; Gonzalez Alam et al., 2022; Jayaprakash et al., 2024; Labache et al., 2023; Liang et al., 2021; Pilli et al., 2019; Wan et al., 2022, 2023). The classical theory of why the cortex exhibits functional asymmetry posits that it avoids costly duplication of neural circuits with the same function (Levy, 1977), which forms the dominant hemisphere or lateralization (Karolis et al., 2019). However, this prediction has been difficult to test because of lacking direct measurements for energy cost across the whole cortex. Here we test this prediction using our energy-gradient model, under the assumption that the spatial pattern of asymmetry in functional brain organization (Gonzalez Alam et al., 2022; Labache et al., 2023; Liang et al., 2021; Wan et al., 2022, 2023) relates to asymmetry in glucose metabolism.

## Results

### Maps of functional organization gradients and glucose metabolism

We first downloaded the FDG-PET and fMRI images for 20 individuals from an open data source (https://openneuro.org/datasets/ds004513/versions/1.0.4). The data description and brain scanning parameters can be seen in the previous publication (Castrillon et al., 2023). In short, fMRI was simultaneously performed with quantitative FDG-PET at the Technical University of Munich with 9 individuals as experimental datasets (4 females; age: 43 ± 7 years) and 11 as replication dataset (6 females; age: 27 ± 5 years). We analyzed the CMRglc map and functional connectome summarized into 360 parcels by a multimodal parcellation scheme (Glasser et al., 2016).

Group-level functional organization gradients were computed by reducing the dimensionality of the affinity matrix of the mean functional connectome across individuals using diffusion map embedding (**Figure 2A**). We computed the 100 principal gradients for 10 sparsity parameters each by thresholding the top 10% (sparsity = 0.9) to the full (sparsity = 0) functional connectome. The sparser the matrix, the higher the variance explained by the gradient framework. For brain gradient maps with sparsity = 0.9, the standard setting of previous works (Dong et al., 2021; Hong et al., 2019; Margulies et al., 2016; Valk et al., 2022; Wan et al., 2022, 2023), G1 separated association areas from the visual cortex, G2 separated somatomotor cortex from the visual cortex, and G3 separated task-positive from task-negative networks. The G4-20 brain maps are shown in **Supplementary Figure S1**. The more similar the gradient loadings of different regions, the more similar the functional connectivity profile of those regions. For gradients 4 to 100, the residuals gradually decrease and include more subtle topographic variance. There is no intercorrelation among these gradients (**Supplementary Figure S2**), indicating the statistical independence of gradients. The group-level CMRglc map **(Figure 2B**) was calculated by averaging the individual CMRglc maps and then z-scoring both CMRglc and gradient maps. This suggests spatial differentiation of glucose uptake, with one anchor in precuneus, and the other anchor in temporal pole regions.

**Figure 2.**
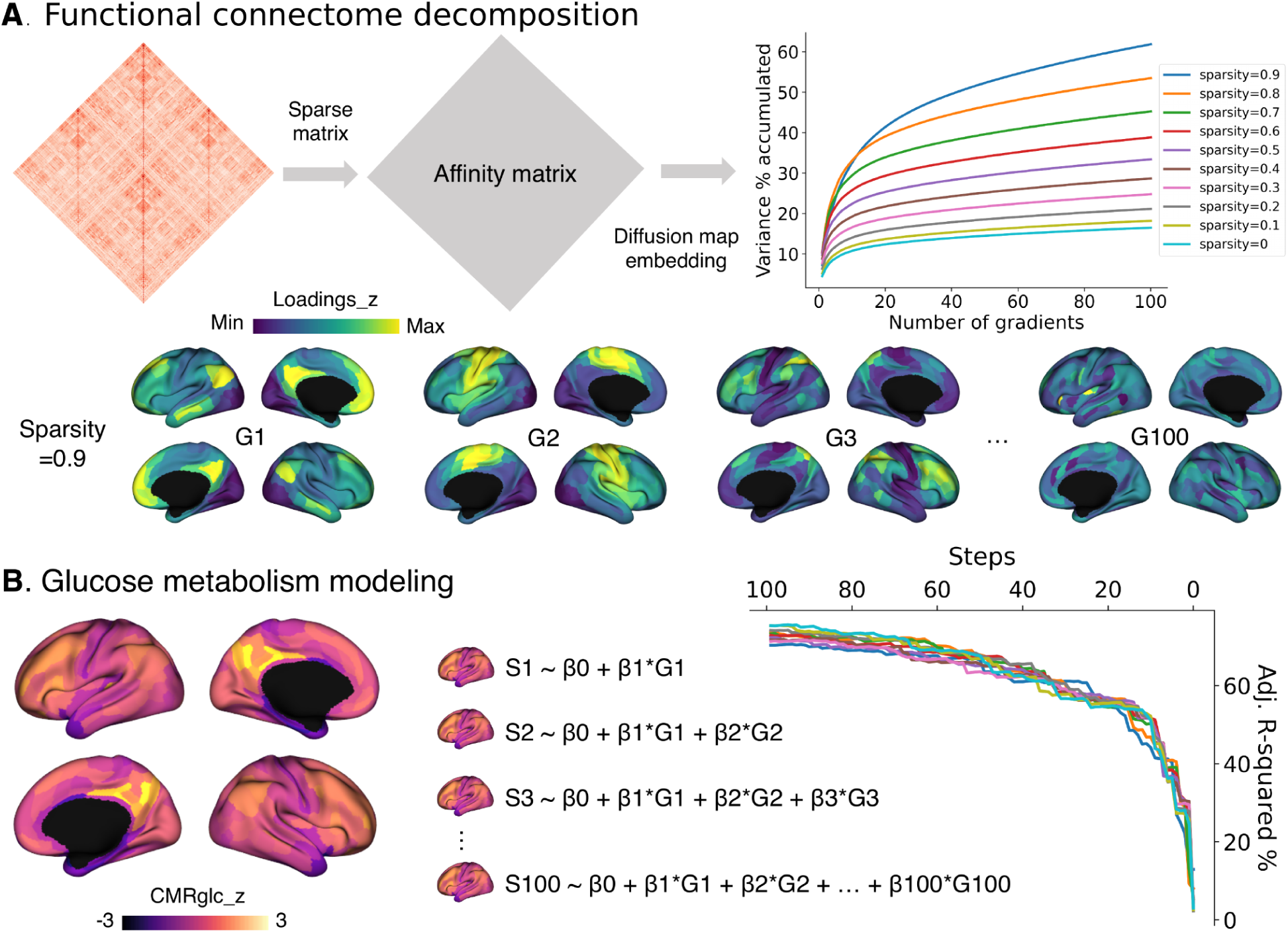
Generating functional organization gradients and glucose metabolism maps. **A)**. illustrates the decomposition processing for the group-level functional connectome. Each sparsity parameter means the threshold of the functional connectome, for example, the top 10% uses sparsity = 0.9 and the full connectome uses sparsity = 0. **B).** shows the group-level CMRglc z-scored map and how the stepwise regression models are generated. The chart displays the adjusted R-squared values of the model by entering more numbers of gradients.

We then generated the stepwise linear regression models without removing variables, as shown in **Figure 2B**. Step1: CMRglc = β0 + β1*G1, step2: CMRglc = β0 + β1*G1 + β2*G2, and so on until step100: CMRglc = β0 + β1*G1 + … + β100*G100. We plotted how the adjusted R-squared values would increase or decrease as more gradients entered the model. It showed that the adjusted R-squared increased logarithmically with the number of gradients. For the last step (step 100), the regression model explained over 70% of the variance of the CMRglc map (sparsity ranging from 0 to 0.9: 75.7%, 75.7%, 74.4%, 71.6%, 72.0%, 72.5%, 73.1%, 73.7%, 73.8%, 70.4%). Moreover, the regression model could explain over 50% of the variance of CMRglc by entering the first 10 gradients (sparsity from 0 to 0.7: 51.1%, 52.2%, 52.0%, 52.0%, 51.8%, 51.5%, 51.7%, 50.3%). However, at sparsity 0.8 and 0.9, 15 (52.6%) and 16 (52.3%) gradients were required to explain more than 50% of the variance, respectively. Together, these results suggest that the way the brain’s intrinsic functional networks are spatially organized is directly related to how much energy a given region uses, that is regions with similar connectivity profiles also consume similar amounts of energy, extending previous notions of connectivity and energy expenditure to a spatial framework of brain organization.

### Model explanation and regularization

Next, to determine whether the regional variance of FC and CMRglc is symmetrically explained by the gradients, that is, whether x amount of variance explained in FC by the first y gradients corresponds to x variance explained in the CMRglc map, we used slope and mean absolute error (MAE) to describe the symmetry and information loss across each sparsity parameter (**Figure 3A**). Slopes close to 1 indicate a full symmetric explanation of gradients to sparse FC matrix and CMRglc map and smaller MAE values indicate less information loss during this procedure. Slopes varied widely across sparsity parameters (from 0 to 0.9): 5.68, 5.16, 4.38, 3.48, 3.04, 2.51, 2.20, 1.93, 1.53, 1.07. Sparsity = 0.9 showed the lowest MAE (sparsity from 0 to 0.9, MAE = 0.248, 0.232, 0.214, 0.169, 0.145, 0.116, 0.092, 0.064, 0.036, 0.013). This indicates that the sparsest functional connectome can explain the CMRglc map most symmetrically, but that for the least sparse gradient, the variance in energy consumption was explained five times better than the variance in the functional connectome. Given that using sparsity = 0.9 made the best symmetry in explaining both functional connectome and CMRglc map, the following analyses were performed for sparsity = 0.9.

**Figure 3.**
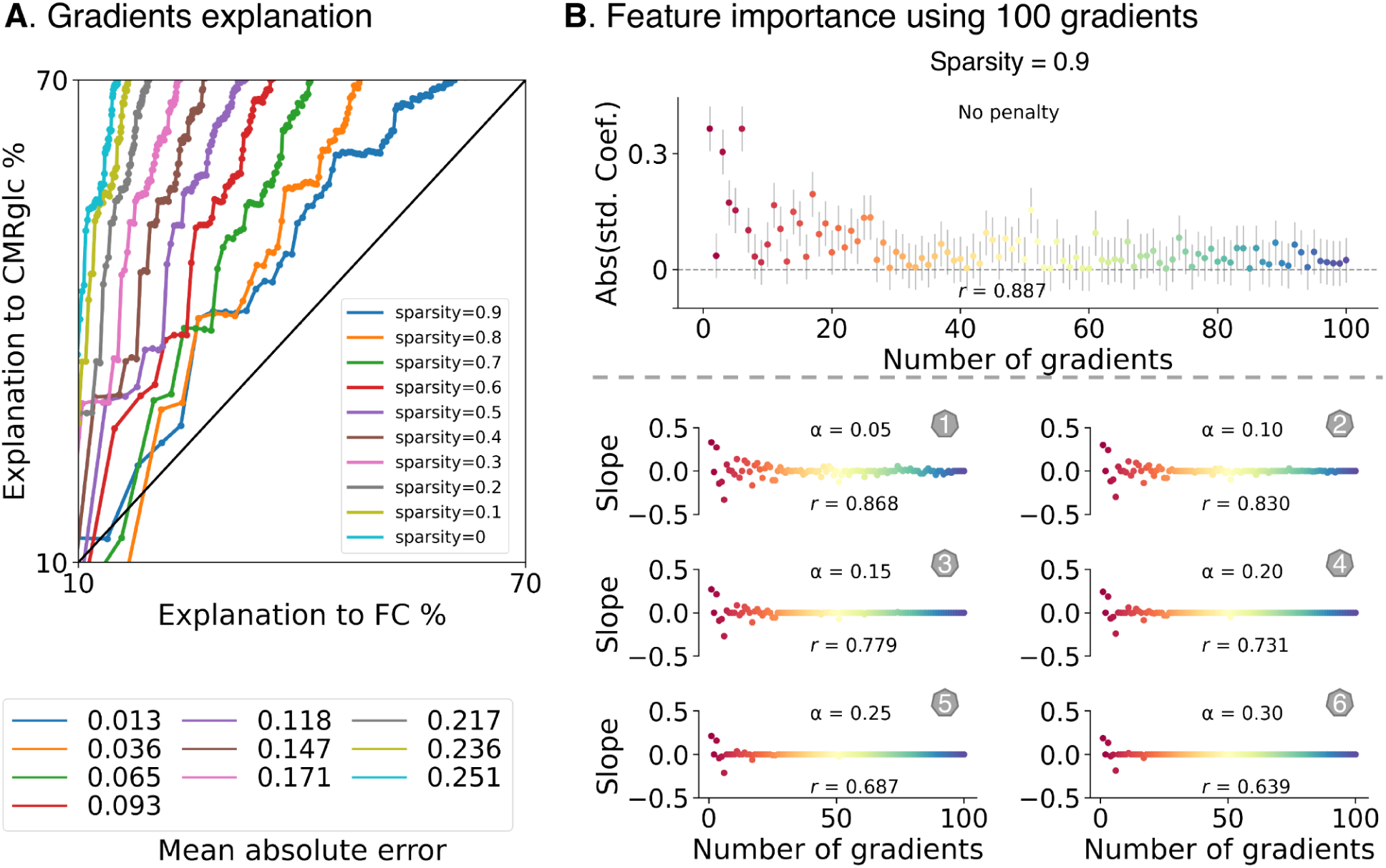
Model explanation and regularization. **A)**. plots the scatter with the x-axis showing FC variance explained by gradients and the y-axis showing CMRglc variance explained by gradients. The lower panel further displays how their relationship (slope) changes along models with entering more gradients (starting from 10). **B).** plots scatter using elastic net regularization (lasso/ridge = 1) onto the step 100 model: one without penalty (standardized regression coefficient and 95% confidence interval by number of gradients), and the other with six penalty parameters (α) from 0.05 to 0.3 (slope by number of gradients). Pearson correlation coefficients (*r*) were calculated using true CMRglc and model-predicted CMRglc maps.

Following this, we used a statistical method called elastic net to simplify our model and test how the different brain gradients are related to energy use in the brain. By adjusting the model’s complexity, we could investigate how much each gradient contributed to predicting brain energy use. We set L1 ratio = 0.5 to balance the lasso and ridge regularization, and manually adjusted penalty parameters (α) from 0.05 to 0.3 with 0.05 intervals for step 100 (**Figure 3B**). We first plotted the absolute values of standardized regression coefficients with a 95% confidence interval (CI) to reveal the relative contribution of each gradient to the fit without including a penalty. The signed values (i.e.. non-absolute) can be seen in **Supplementary Figure S3.** The top five slopes were: G1 [0.365(0.308, 0.421)], G6 [-0.365 (-0.421, -0.308)], G3 [0.305 (0.248, 0.361)], G17 [-0.195 (-0.252, -0.139)], and G4 [-0.173 (-0.230, -0.117)] with the association between true CMRglc and model predicted CMRglc to be 0.887 (Pearson *r*). With increasing penalty parameters, the Pearson *r* between true CMRglc and model-predicted CMRglc was reduced and first gradients were still contributable, indicating that the order of gradient may follow the energy consumption principle, i.e., accumulative gradients by the order. Overall we found that the first few gradients had the strongest influence on the energy expenditure map, with one gradient (G17) explaining more variance than expected, and G2 less. This illustrates that low dimensions that explain a lot of variance in intrinsic functional dimensions also explain a lot of regional variance in energy maps - further underscoring the link between functional topology and brain metabolism.

### Validation using null models

We further tested whether the gradient maps (sparsity = 0.9) could predict a surrogate CMRglc map of random distribution generated by variogram (**Figure 4A**). We permuted the CMRglc map 1000 times using spatial autocorrelation (Burt et al., 2020; Leech et al., 2023; Shinn et al., 2023; Vos de Wael et al., 2020), and obtained 1000 spatially autocorrelated CMRglc surrogate maps based on the geometric distance matrix. We observed that the true model exceeded 96.5% of the variogram autocorrelated models. We also permuted the gradient maps 1000 times using the variogram method, and it showed that the true model exceeded 100% of the variogram models (**Figure 4B**). This indicates that the gradient-energy relationship is generated by the ‘true’ CMRglc map but not by random spatial patterns, providing further validity for our observations.

**Figure 4.**
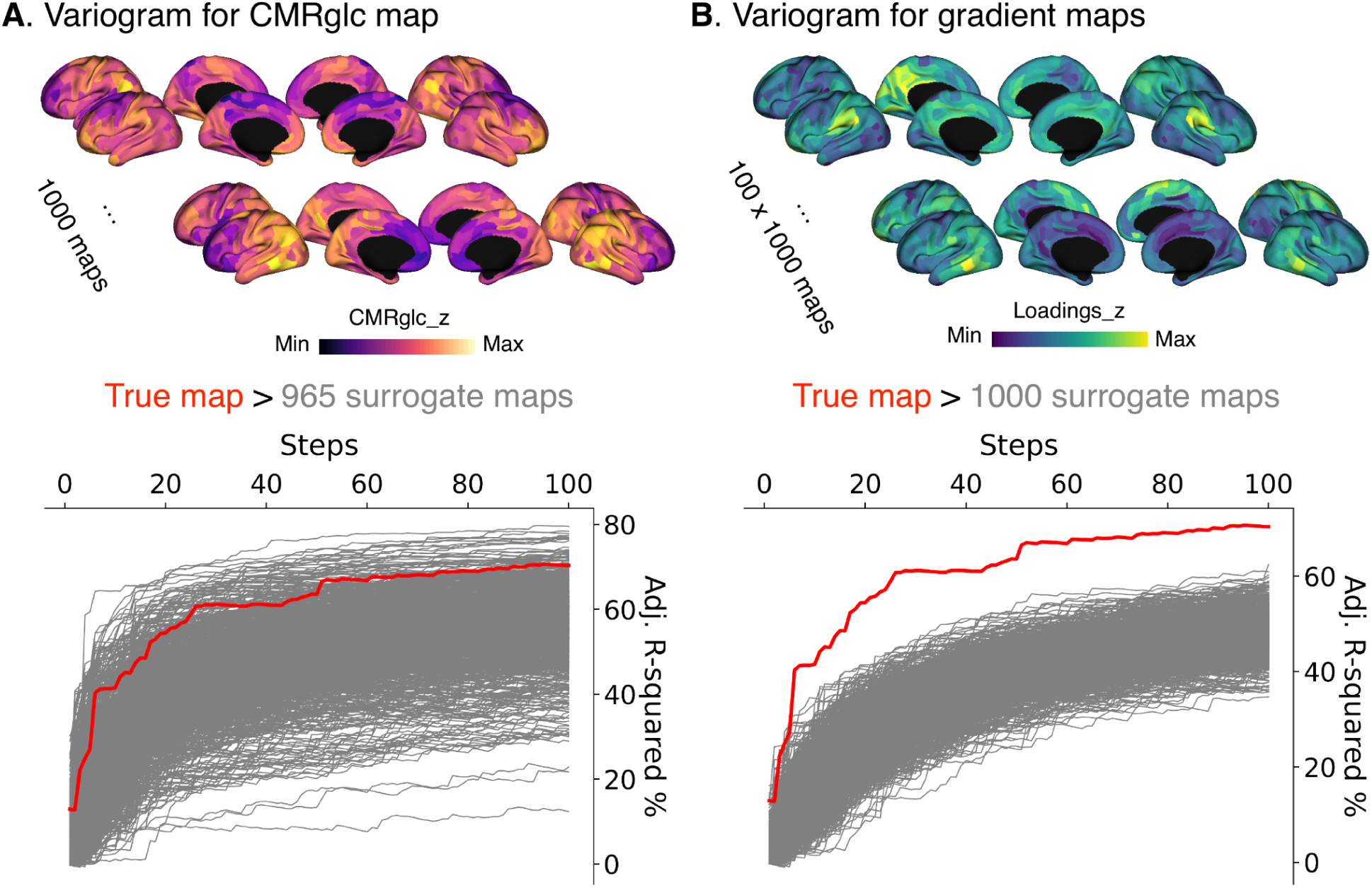
Null models autocorrelated surrogate maps (sparsity = 0.9). **A)** and **B)** show 1000 surrogate maps for CMRglc map and gradient maps using variogram spatial autocorrelation based on the geometric distance matrix. Red and gray lines indicate true and null models.

### Asymmetry modeling

Following we wished to apply our model of energy consumption and brain organization to the asymmetry of brain organization, motivated by the hypothesis stating that brain asymmetry optimizes brain energy consumption. To do so, we calculated hemispheric gradients in the left and right functional connectome separately and aligned the right hemisphere (RH) gradients to the left hemisphere (LH) gradients using Procrustes rotations (**Figure 5A**), finally calculated LH-RH for each gradient as functional asymmetry maps, as in our previous work (Wan et al., 2022, 2023). The CMRglc asymmetry map was calculated by z-scoring the left (LH) and right hemispheres (RH) separately (**Figure 5B**). Our gradient asymmetry maps were similar to those reported in previous studies (Gonzalez Alam et al., 2022; Labache et al., 2023; Liang et al., 2021; Wan et al., 2022, 2023). We observed that there was a strong correlation (*r* = 0.732 with 100 gradients) but no improvement in the model performance curve (adjusted R-squared maximum: 2.3% with 23 gradients, **Figure 5C**), indicating that a highly complex model of many gradients is needed to explain metabolic asymmetry.

**Figure 5.**
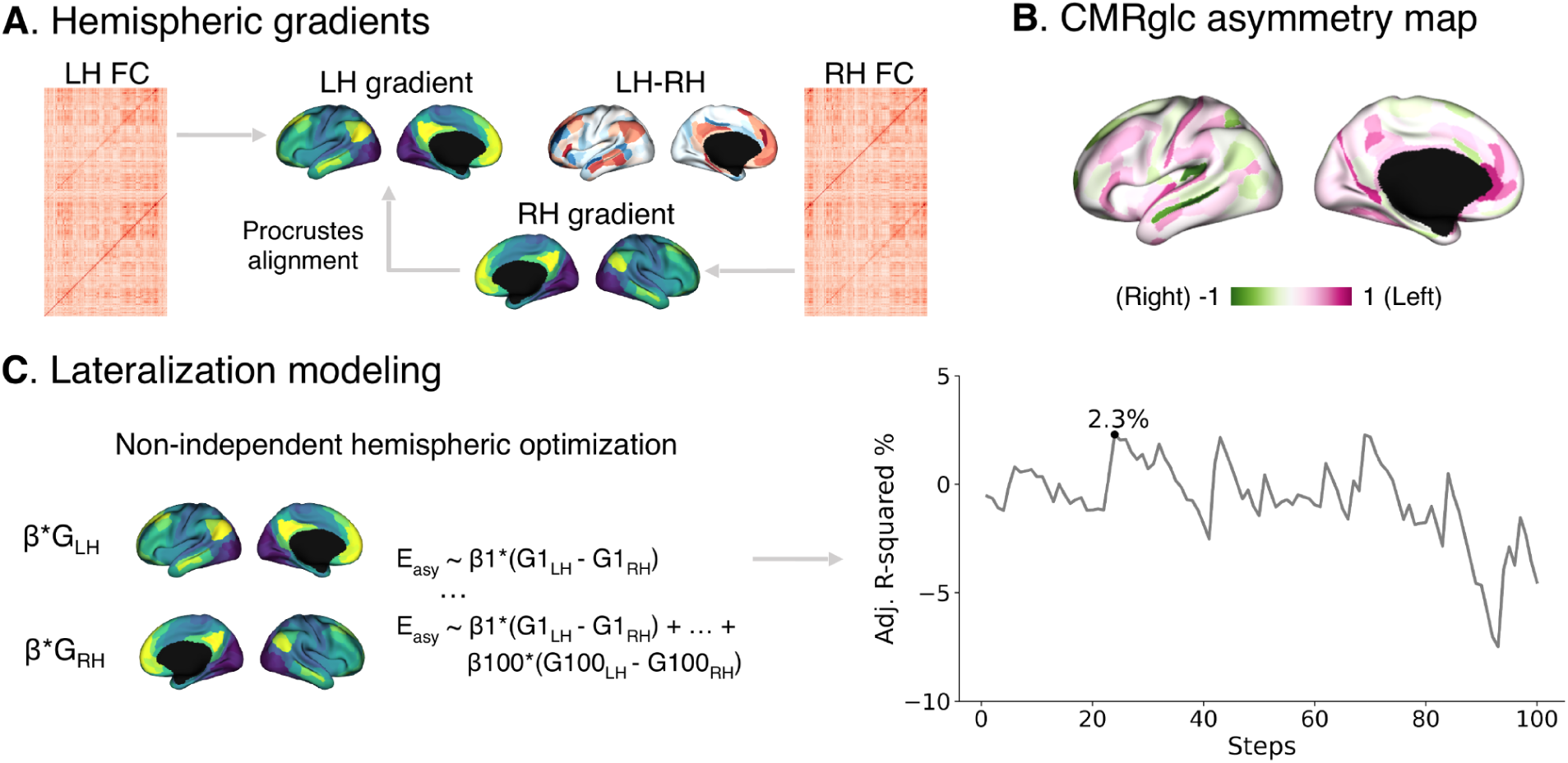
Asymmetry modeling. **A)**. illustrates how hemispheric gradients are calculated and an example asymmetry map of G1. **B).** shows the CMRglc asymmetry map. **C)**. visualizes the lateralization modeling with non-independent hemispheric optimization for energy and how adjusted R-adjusted values change with regression steps.

To further investigate this observation, we used elastic net regularization to increase the penalty and Pearson *r* to indicate the model fit instead of reporting adjusted R-squared. Increased α (penalty parameter on the complexity of model) from 0.05 to 0.3, decreased model fit (Pearson *r*) from 0.677 to 0.293, and there was no trend for which gradients were zeroed, e.g., pre- or post-sequential gradients (**Supplementary Figure S5**). We further evaluated collinearity as a possible reason for lacking increased fit with the increased number of gradients, by correlating the gradient asymmetry maps. Yet we found few collinearities (**Supplementary Figure S6**). Together, these observations indicate that asymmetric patterns in energy consumption may not rely on the explained variance of functional organization on its topology (as was the case for the entire cortex modeling in **Figure 2B**, where the order of gradients was found to be important).

In addition, we also aligned the LH to RH gradients to test whether the results could be affected by inter-hemispheric alignment (**Supplementary Figure S4-6**). The gradient asymmetry maps were similar between the two alignment approaches, and the regularization for model step 100 showed that Pearson r decreased from 0.662 to 0.307 (**Supplementary Figure S5**). The maximum adjusted R-squared reached 7.6% with 51 gradients **(Supplementary Figure S7**).

### Hemispheric independence

Though we found that the variance in energy asymmetry can be explained by the combined asymmetry of gradients, this relationship does not follow a clear gradient order, which contradicts the principle of accumulative brain functions with each additional gradient. To address this, we conducted additional analyses—first by modeling each hemisphere separately, and then by generating energy-guided gradient asymmetry maps. These steps were designed to answer a key question: does considering the hemispheres as independent organizational systems provide a better explanation of energy use than considering them as interdependent? (**Figure 6A**). To answer this question, we trained models on the LH data and then used the trained parameters to predict CMRglc in the RH and vice versa for another hemisphere. We observed that each hemisphere performed well on its own, but slightly worse when predicting the other hemisphere. Similar results were obtained using different hemispheres for alignment (**Supplementary Figure S8A**).

**Figure 6.**
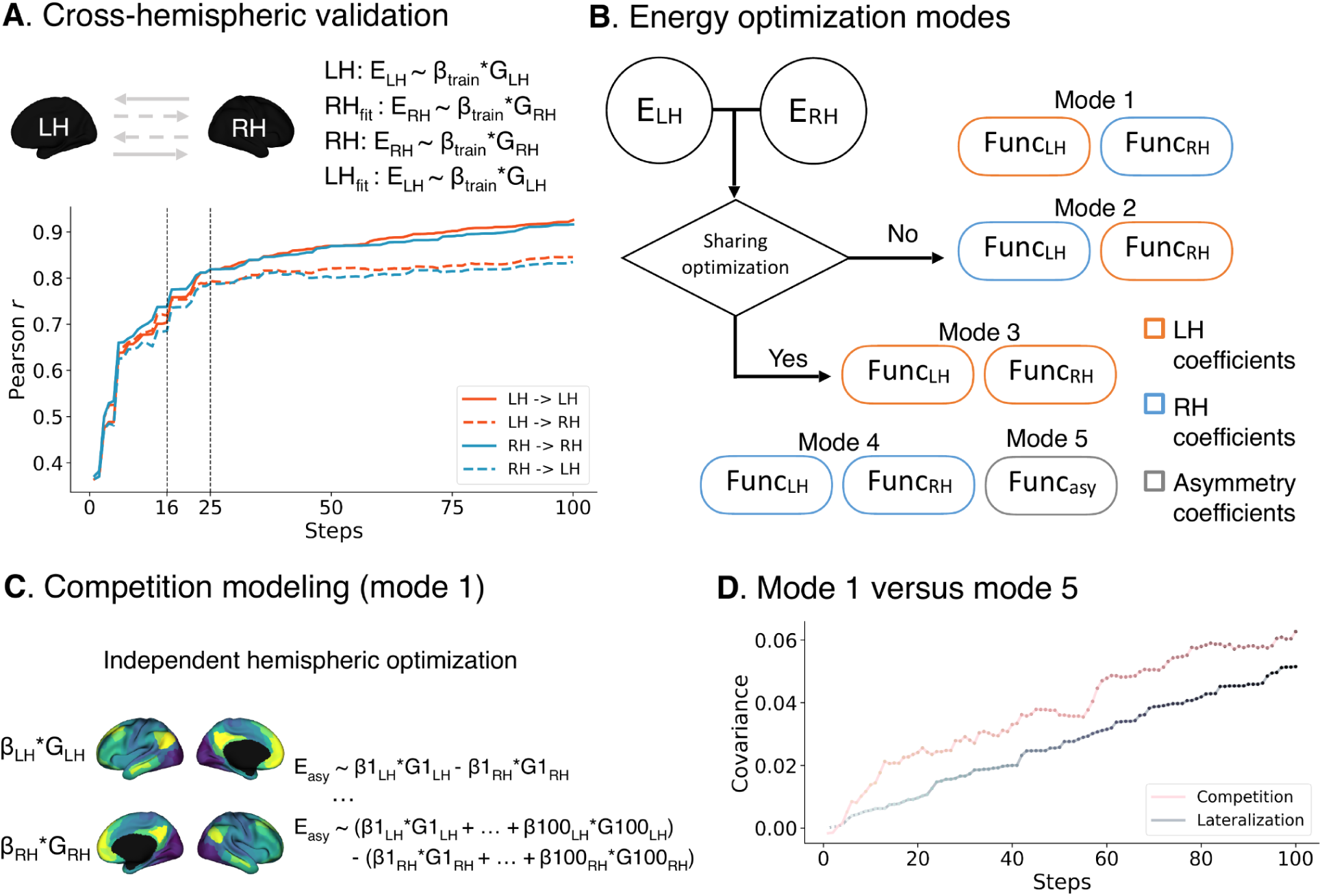
Hemispheric modeling. **A)**. visualizes the cross-hemispheric validation for the models and performance in the training and fitting sets. **B)**. illustrates five optimization modes depending on whether LH and RH share the energy-gradient coefficients. If not, modes 1 and 2 use independent LH and RH training coefficients themselves (mode 1) or exchange (mode 2). If yes, modes 3, 4, and 5 share the LH (mode 3) or RH (mode 4) or asymmetry (mode 5) coefficients. **C)**. visualizes how to calculate hemispheric differences after the modes above, e.g., the competition modeling (mode 1). **D).** compares fitting covariance between competition (mode 1) and lateralization (mode 5). All mode comparisons can be seen in **Supplementary Figure S3**.

To further determine whether LH and RH have a similar or different energy map-functional organization relationship, we set up five different ‘energy sharing’ models (**Figure 6B**). If LH and RH have a differentiable association between energy map and functional organization, LH and RH use independent training coefficients themselves (mode 1) or in exchange (mode 2). If they do, both hemispheres would share the LH (mode 3), RH (mode 4), or asymmetry (mode 5) coefficients. We defined the asymmetry-based model representative for ‘hemispheric lateralization’ (**Figure 5C**, reflected by mode 5) and the mode that modeled LH and RH separately for ‘hemispheric competition’ (**Figure 6C**, mode 1). Instead, modes 2, 3, and 4 were using the opposite hemispheric coefficients. Then, we compared the best competition and lateralization modes between competition and lateralization (**Figure 6D**). We used covariance instead of correlation to indicate the closeness between predicted and true energy asymmetry maps to constrain both within one scale. It showed covariance scores in the competition were always higher than lateralization (step 100 mode comparison ratio = 1.21:1), reflecting hemispheric competition (e.g., two separate models for LH and RH) may better explain CMRglc asymmetry. This result was consistent using different hemispheres for alignment (**Supplementary Figure S8B**).

Other mode comparisons are shown in **Supplementary Figure S9**. The covariance scores between true and predicted CMRglc asymmetry maps for these modes in step 100 were -0.030 (mode 2: competition), 0.020 (mode 3: lateralization), and 0.013 (mode 4: lateralization). This suggests the competition model performs better than the lateralization model even with cross-hemispheric optimizing parameters.

### Validation at vertex and individual levels

Finally, we evaluated the robustness of our group-level results of the entire cortex and asymmetry with downsampled fsLR-5k vertex analyses with symmetric 4428 vertices per hemisphere without midline. We obtained 100 gradients using sparsity = 0, 0.5, and 0.9 (**Supplementary Figures S10-12**). When using 100 gradients in the model, the adjusted R-squared values were 36.0%, 37.4%, and 43.0% (**Supplementary Figure S11**). Regarding the modeling of asymmetry maps (sparsity = 0.9), the adjusted R-squared value was 8.3% for all 100 gradients. However, the penalty parameter (α) had a large effect on the fit of the 100 gradient models (**Supplementary Figure S13**). As α increased from 0.05 to 0.3, model fit (Pearson *r*) decreased from 0.279 to 0. The competition model was always better than the lateralization model (**Supplementary Figure S14**).

Second, to assess whether our observations of the entire cortex and asymmetry extend beyond the group level, we tested the models at the individual level (N = 20). We used the raw individual gradients for the whole cortex modeling and adjusted individual gradients (to the group level) for asymmetry modeling (**Supplementary Figure S15-21**). Overall, our group-level findings were also present at the individual level. Specifically, for the stepwise regression models, the adjusted R-squared values increased as more gradients entered the CMRglc map for all subjects, and the mean explained variance was 44.2% with a standard deviation of 8.3% (**Supplementary Figure S15**). Four individuals’ gradient maps (**Supplementary Figure S16**) and CMRglc (**Supplementary Figure S17**) maps were shown to illustrate the robustness of our model at the level of the individual. The accumulated explained variance of gradient decomposition had comparable levels for all subjects (mean slope = 0.778, standard deviation = 0.171, **Supplementary Figure S18**). Regarding asymmetry modeling, as with the group level, no individual showed an increasing trend for the adjusted R-squared, and the maximum adjusted R-squared was 3.3% with 19 gradients. Regarding the individual cross-hemispheric validation, the mean performance levels of RH-to-LH and LH-to-RH were highly similar, but they showed high variability at the individual level (**Supplementary Figures S20 and 21**). Regarding the comparison between competition and lateralization models, competition performed better than lateralization for 95% of the individuals (**Supplementary Figure S22**).

## Discussion

In this study, we investigated how the functional organization of the brain is related to its energy consumption, focusing specifically on regional energy consumption using PET imaging of glucose metabolism (CMRglc). We hypothesized that the systematic functional organization of the brain is directly related to regional variability in energy consumption. To test this, we used a stepwise linear regression model in which brain energy is predicted by functional gradients representing unique global patterns of brain intrinsic co-activation. Our model explained approximately 70% of the variance in energy consumption at the group level, a robust result that held across different levels of sparsity in the functional connectivity (FC) matrix. Interestingly, gradients generated from sparser FC networks were better at capturing regional variability in energy expenditure, suggesting that the strongest functional connections in each region play a key role in determining how much energy the brain consumes. We also used these models to investigate how differences in energy use between the two hemispheres of the brain relate to their functional organization asymmetry. By comparing models with independent and non-independent optimization for each hemisphere, we found that energy asymmetry was better explained when the hemispheres were treated as separate systems, supporting theories of hemispheric competition. Our findings held when tested at the individual level, further confirming the robustness of our results. Overall, our study reveals an intrinsic link between the spatial organization of intrinsic brain function and its energy expenditure, providing new insights into the energetic basis of brain function and how the brain manages its resources.

First, we examined whether regional variability in energy expenditure can be explained by intrinsic functional organization gradients. By accumulating organizational gradients in our model, we found that five gradients capture 27% of the variance in the energy expenditure map, ten gradients capture 40%, and about 100 gradients capture about 70%. Previous studies have shown moderate correlations between energy intake distribution and brain function, with Pearson *r* values ranging from 0.3 to 0.5 (Castrillon et al., 2023; Riedl et al., 2014; Stiernman et al., 2021; Tomasi et al., 2013; Wang et al., 2021). Our model, which uses the topography of intrinsic brain function, achieves a stronger relationship. Notably, gradients based on the top 10% of the connectivity synchronized strongly with the energy map in terms of variance explained in the functional connectivity matrix and the energy consumption map. Using more (i.e. less sparse) information from the functional connectome did not improve the model fit to the energy map. This suggests that regional energy consumption strategies may most closely link to the strongest connections in the functional connectome and not so much to weaker connectivity patterns.

Conceptually, weak connections may not consume as much energy (reduced impact on glucose metabolism). One possible explanation could be that hub regions consume more energy than non-hub regions (Riedl et al., 2014) and that energy consumption of weak connections is influenced by stronger connections, such as hub regions (Power et al., 2013), in an ‘energy hierarchy’. Indeed, functional hubs serve as connectors between different brain modules or within each module (Heuvel & Sporns, 2013; Power et al., 2013). For example, the precuneus has been identified as a hub region (Heuvel & Sporns, 2013; Power et al., 2013) and is also consistently a high energy expenditure region in our study, also seen in previous research (Palombit et al., 2022; Stiernman et al., 2021; Tomasi et al., 2013). Another factor that may explain the observed patterns may be redundant connections. A non-redundant model of energy expenditure suggests that sensory areas show redundant connectivity, while parietal and frontal areas show non-redundant connectivity (Salvador et al., 2017). This is consistent with the energy expenditure map observed in our study, as parietal and frontal cortices also have high energy expenditure and sensory areas have low energy expenditure. By aggregating the connectivity per region, previous work found a correlation with the energy expenditure map (correlation coefficient around 0.5) (Castrillon et al., 2023). Identifying the principle or threshold of redundant connectivity, in terms of energy expenditure, could improve our understanding and interpretation of the energy expenditure map. For example, different thresholds could be applied to different brain regions. Overall, an energy-driven understanding of brain function may provide novel perspectives on the relationship between computational demand and energy supply in the human brain across the lifespan.

Brain lateralization has been proposed as a mechanism for optimizing brain energy requirements (Levy, 1977). Here we evaluated this hypothesis by using asymmetry maps of functional organization to model the energy asymmetry map. We observed that the asymmetry model was overfitting. This could not be explained by the possibility of variable independence as only a few gradient asymmetry maps were correlated with each other. This suggests that energy asymmetry may not result solely from cumulative functional lateralization in topology. While asymmetry of functional gradients has been extensively studied (Gonzalez Alam et al., 2022; Labache et al., 2023; Liang et al., 2021; Wan et al., 2022, 2023), our results suggest that gradients may not only reflect static brain organization but also how the brain operates across different spatial scales or in the form of spatial competition in the context of energy optimization. Such an interpretation could be in line with conceptualizations that functional lateralization results from interactions between hemispheric functions (Gotts et al., 2013; Hartwigsen et al., 2021). For example, hemispheric competition describing the changing role of hemispheres has been evidenced during attention and sleep over time (Cohen et al., 1994; Fenk et al., 2023; Kinsbourne, 1977, 1993). Energy lateralization may be the result of such functional spatial competition.

To sum up, here we revealed how cortical energy expenditure is related to spatial maps of functional brain organization. In doing so, we provide a novel framework for functional organization in the context of its energy landscape, illustrating how the topological organization of intrinsic function may relate to the optimization of metabolic processes. These insights may hold relevance for a basic understanding of the biological basis of intrinsic functional activity in the human brain, and provide further perspectives for its changes in neuropsychiatric and neurological disorders.

## Methods

### Data source

We reanalyzed the data from OpenNeuro (Data Number: ds004513). Briefly, data collection was conducted at the Technical University of Munich with 9 individuals as experimental datasets (4 females; age: 43 ± 7 years) and 11 as replication datasets (6 females; age: 27 ± 5 years). The FDG-PET scans contain 5 frames * 300 s/frame. CMRglc map was calculated by multiplying the net uptake rate constant (*K*_i_) based on the 5 frames after motion-corrected, smoothed (FWHM = 6mm), and partial volume corrected using the gray matter (GM), white matter (WM), and cerebrospinal fluid (CSF) masks derived from the T1 images. Finally, the individual CMRglc map was registered to MNI152NLin6ASym 3-mm template and volume to surface mapped to mid-thickness of fsLR-32k space and downsampled to fsLR-5k space and summarized into 360 parcels using multimodal parcellation (Glasser et al., 2016). The fMRI scans contain 300 time series (repetition time = 2000 ms). The images were motion-corrected, high-pass (0.01-0.1 Hz) filtered, regressed out GM, WM, and CSF signals, and finally registered to the native surface spaces, then registered to fsLR-32k space (FWHM = 10mm) and downsampled to fsLR-5k space. The first 5 time points were dropped and then the 295 time series of signal-to-noise ratio (SNR) were summarized into 360 parcels using MMP. The functional connectome of each individual was calculated by the Fisher-z transforming the time series correlation matrix. It was computed using a toolbox MICAPIPE (Cruces et al., 2022) (https://micapipe.readthedocs.io/en/latest/). Detailed data description and image parameters can be seen in the previous publication (Castrillon et al., 2023).

### Gradients of functional organization

After obtaining the individual functional connectome, we calculated the affinity matrix in a normalized angle with different sparsity parameters from 0 (full connectome) to 0.9 (top 10% connectivity of the connectome). Next, we employed the nonlinear dimensionality reduction technique to generate 100 principal gradients for each individual and at the group level. We then set the group-level gradients as the template and aligned each individual gradient with Procrustes rotation to the template. Finally, the comparative individual functional gradients were assessed. Gradients reflect the eigenvectors and gradient loadings reflect the eigenvalues. Cortical regions that are strongly interconnected, by either many suprathreshold edges or a few very strong edges, are closer together. Nodes with little connectivity similarly are farther apart. Regions having similar connectivity profiles are embedded together along the eigenvector. All steps were accomplished in the Python package Brainspace (Vos de Wael et al., 2020). The name of dimensionality reduction, which belongs to the family of graph Laplacians, is derived from the equivalence of the Euclidean distance between points in the diffusion-embedded mapping (Coifman et al., 2005; Margulies et al., 2016; Vos de Wael et al., 2020). It is controlled by a single parameter α, which controls the influence of the density of sampling points on the manifold (α=0, maximal influence; α=1, no influence). Based on the previous work (Margulies et al., 2016), we followed recommendations and set α=0.5, a choice that retains the global relations between data points in the embedded space and has been suggested to be relatively robust to noise in the covariance matrix.

### Stepwise regression models

We then generate the stepwise regression models to observe how the explanation of variance (adjusted R-squared) changes along the steps. We set the CMRglc map as a dependent variable and every gradient as an independent variable. Ordinary forward regression kicks out the insignificant independent variables during stepwise model generation. Here, we don’t kick out any gradients and put the gradient maps to the model by eigenvector order (i.e., the number of gradients shown in the figures). Ordinary least squares (OLS) estimation in the Python package statsmodels (https://www.statsmodels.org/stable/index.html) was used to generate the models. We extracted 100 gradients so there were 100 steps or regression models finally. Adjusted R-squared is a modified version of R-squared that has been adjusted for the number of independent variables in the model. Adjusted R-squared = 1 - [SS_residual_ / (n-k)] / [SS_total_ / (n-1)], where n indicates brain data point (i.e., 360 for MMP and 8856 for fsLR-5k) and k indicates the number of gradients in the model. Adjusted R-squared decreases or is negative when adding more gradients.

In addition, we regularized the step 100 model using balanced lasso and ridge algorithms. We manually tuned the penalty parameter from 0.00005 to 0.0003 to visualize which gradients can be regularized and how much it can influence the model fitting.

### Asymmetry

To quantify the inter-hemispheric differences, we calculated asymmetry by LH minus RH. Regarding the CMRglc, we z-scored LH and RH separately and then calculated the energy uptake asymmetry map. Regarding gradients, we computed the LH and RH gradients, then aligned RH to LH gradients to make them comparable in the main figures. In addition, we also aligned LH to RH gradients to test the alignment robustness, shown in supplementary figures. Finally, we could calculate the gradient asymmetry maps.

### Code availability

All analyses were conducted based on Python 3.8 and the main dependencies are BrainSpace (https://brainspace.readthedocs.io/en/latest/index.html) and statsmodels (https://www.statsmodels.org/stable/index.html). Scripts for this study are shared at a GitHub repository (https://github.com/wanb-psych/asymmetry_energy).

### Data availability

All raw and preprocessed resting state fMRI and PET data are available through OpenNeuro, which can be downloaded at https://openneuro.org/datasets/ds004513/versions/1.0.4. In addition, the group-level and individual-level FC matrix and CMRglc maps are available at the above GitHub repository, which ensures reproducible results with the scripts.

## Acknowledgments

The idea was inspired by the geometric constraints on brain function paper (Pang et al., 2023). BW is supported by the International Max Planck Research School on Neuroscience of Communication: Function, Structure, and Plasticity (IMPRS NeuroCom). VR is supported by the European Research Council (ERC) under the European Union’s Horizon 2020 research and innovation program (ERC Starting Grant, ID 759659). SLV is funded by the Otto Hahn Award at the Max Planck Society and Helmholtz International BigBrain Analytics and Learning Laboratory (HIBALL), supported by the Helmholtz Association’s Initiative and Networking Fund and the Healthy Brains, Healthy Lives initiative at McGill University.

## Author contributions

Bin Wan: Conceptualization, Methodology, Formal analysis, Writing - Original Draft, Writing - Review & Editing, Visualization, Project administration, Funding acquisition.

Valentin Riedl: Data Collection, Writing - Review & Editing

Gabriel Castrillon: Data Collection, Writing - Review & Editing

Matthias Kirschner: Writing - Review & Editing.

Sofie L. Valk: Writing - Original Draft, Writing - Review & Editing, Supervision, Funding acquisition.

## Supplementary information

Supplementary Figures 1–22.

---

**Figure S1.**
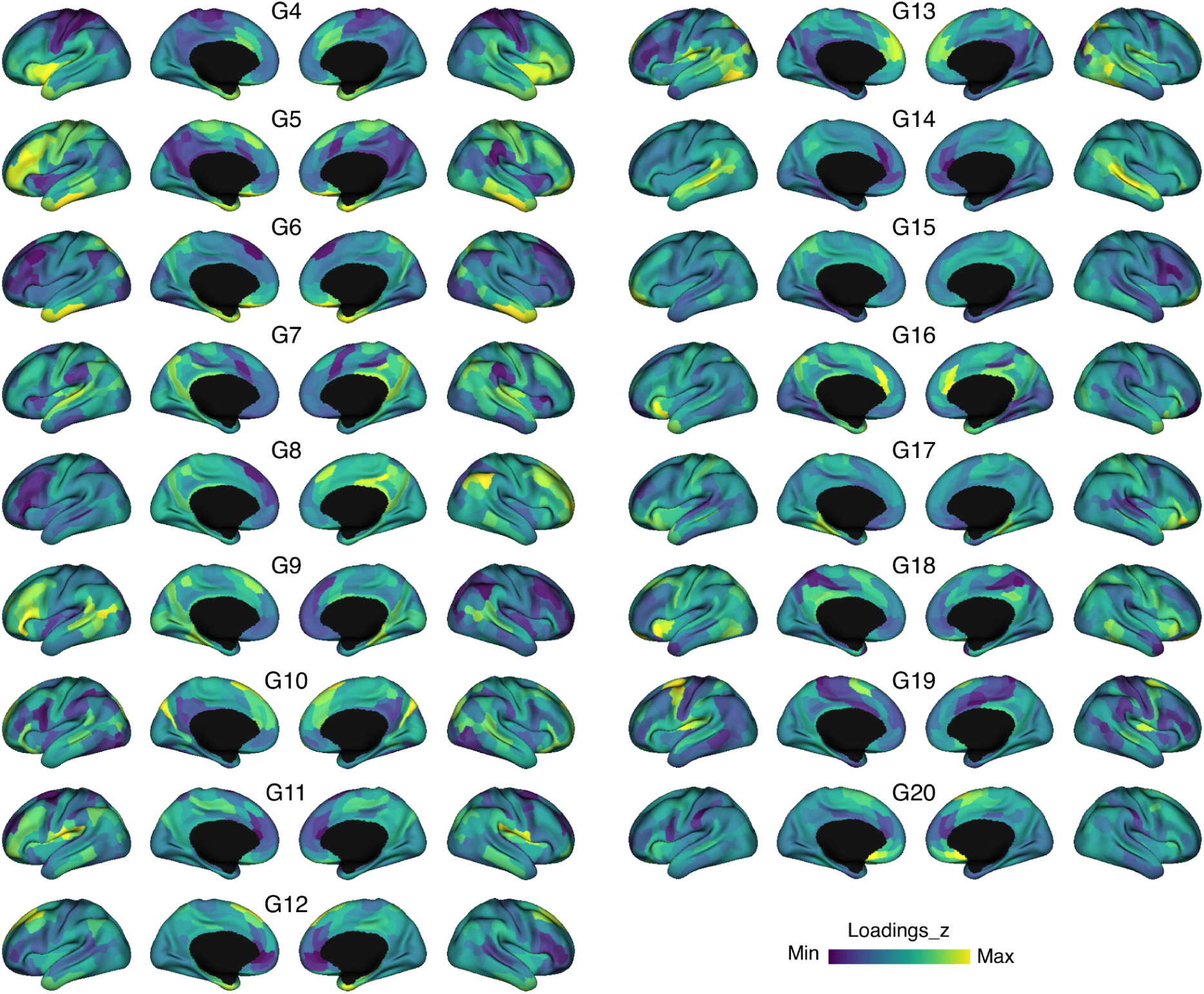
G4-20 of the group-level functional connectome.

**Figure S2.**
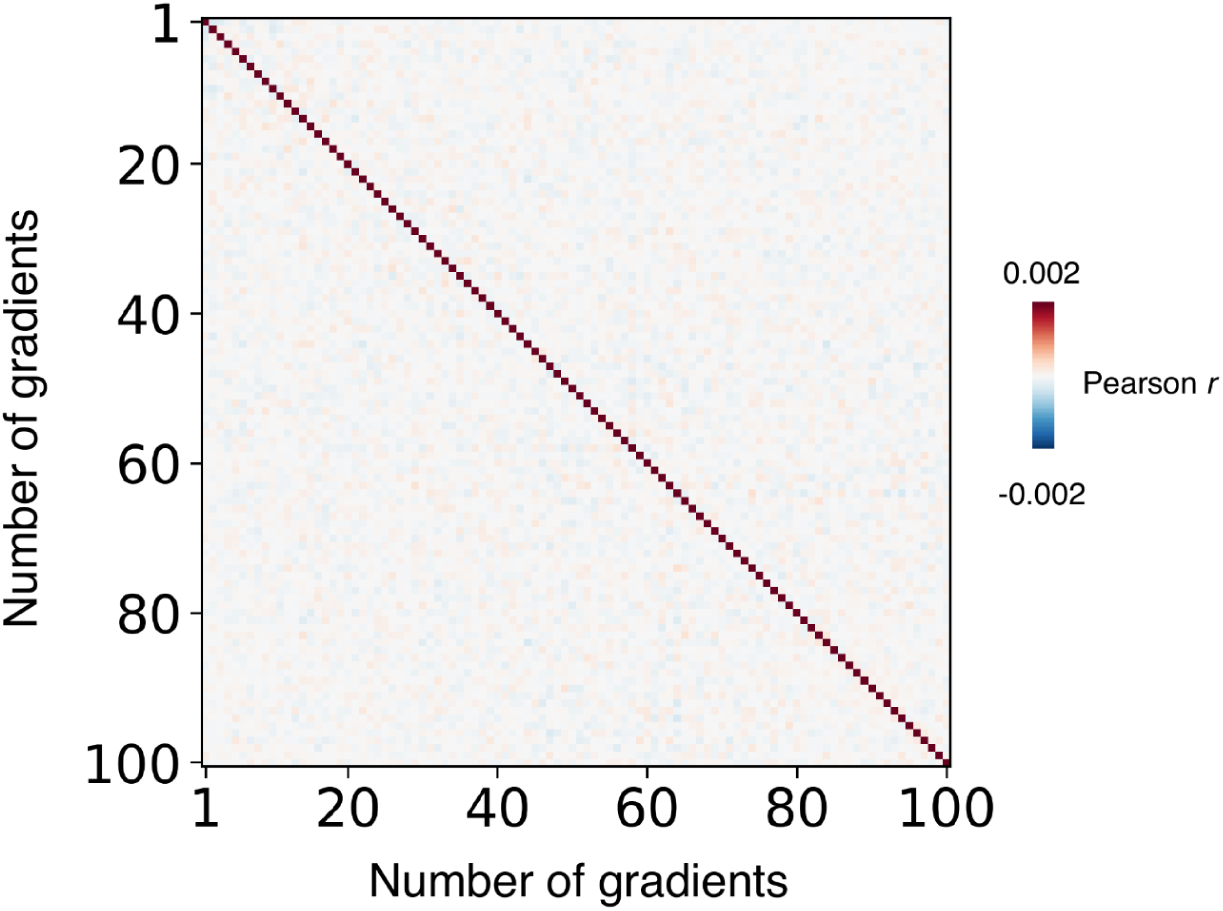
Inter-gradient correlation to identify the independence of each gradient.

**Figure S3.**
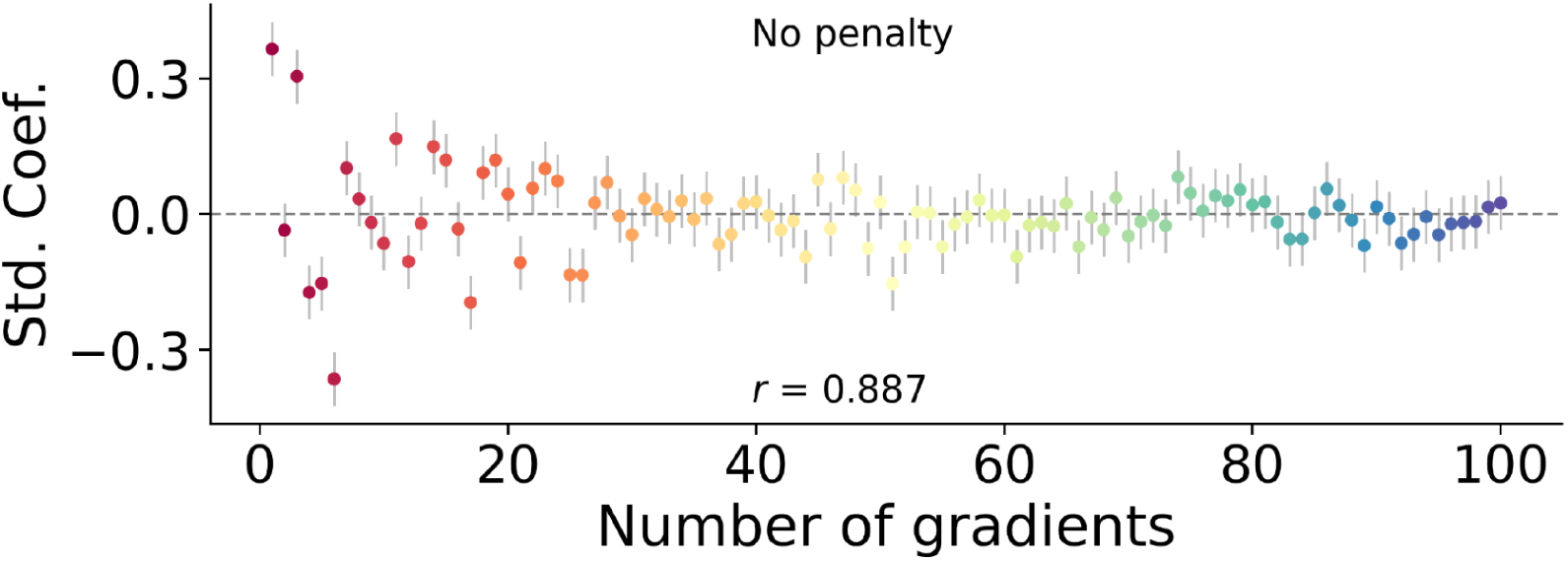
Standardized regression coefficient and 95% confidence interval for non-penalty model.

**Figure S4.**
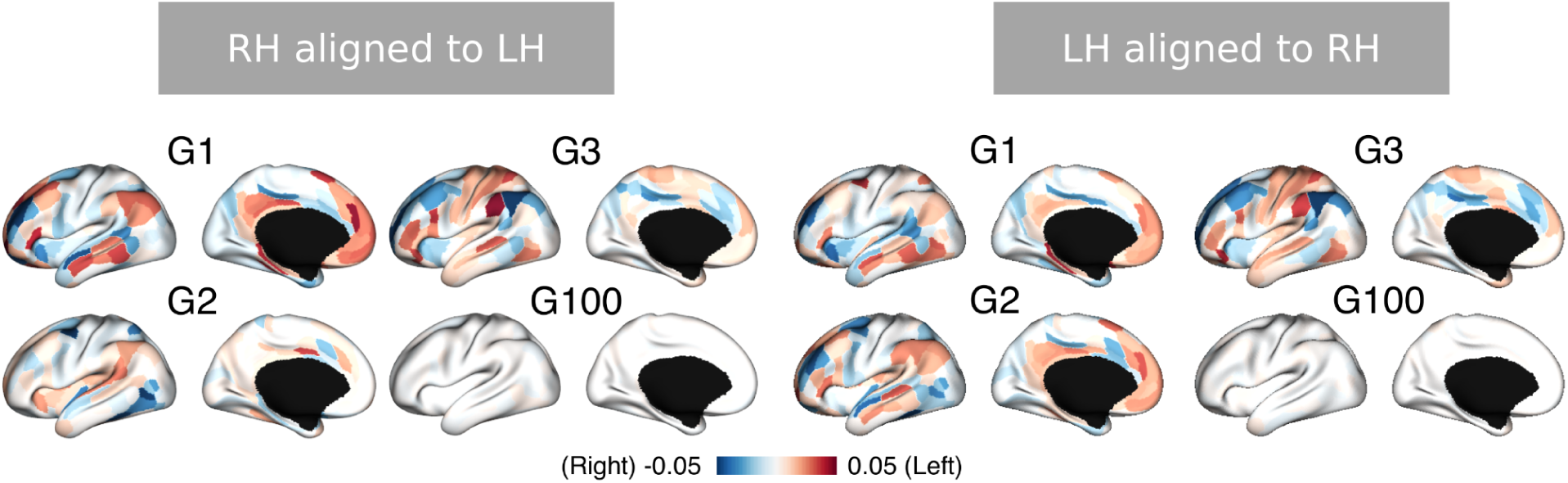
Gradient asymmetry maps for different alignment template references. Left and right panels are LH and RH as the templates, respectively.

**Figure S5.**
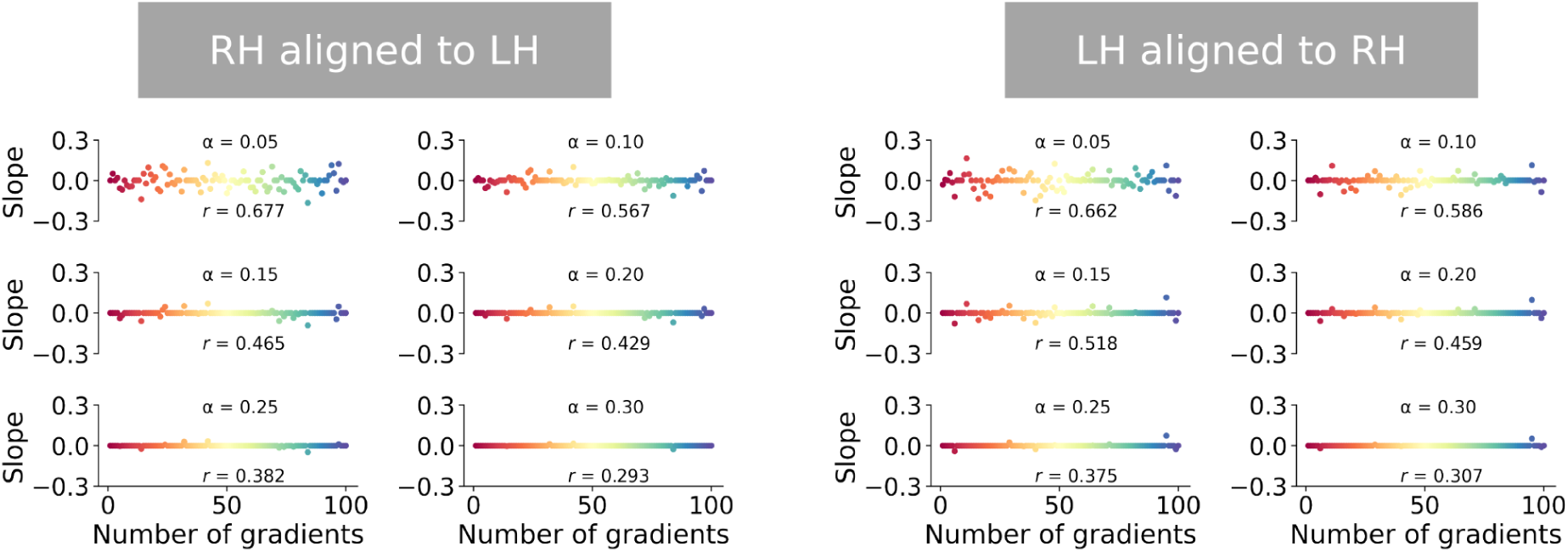
Model regularization for different alignment template references. Left and right panels are LH and RH as the templates, respectively.

**Figure S6.**
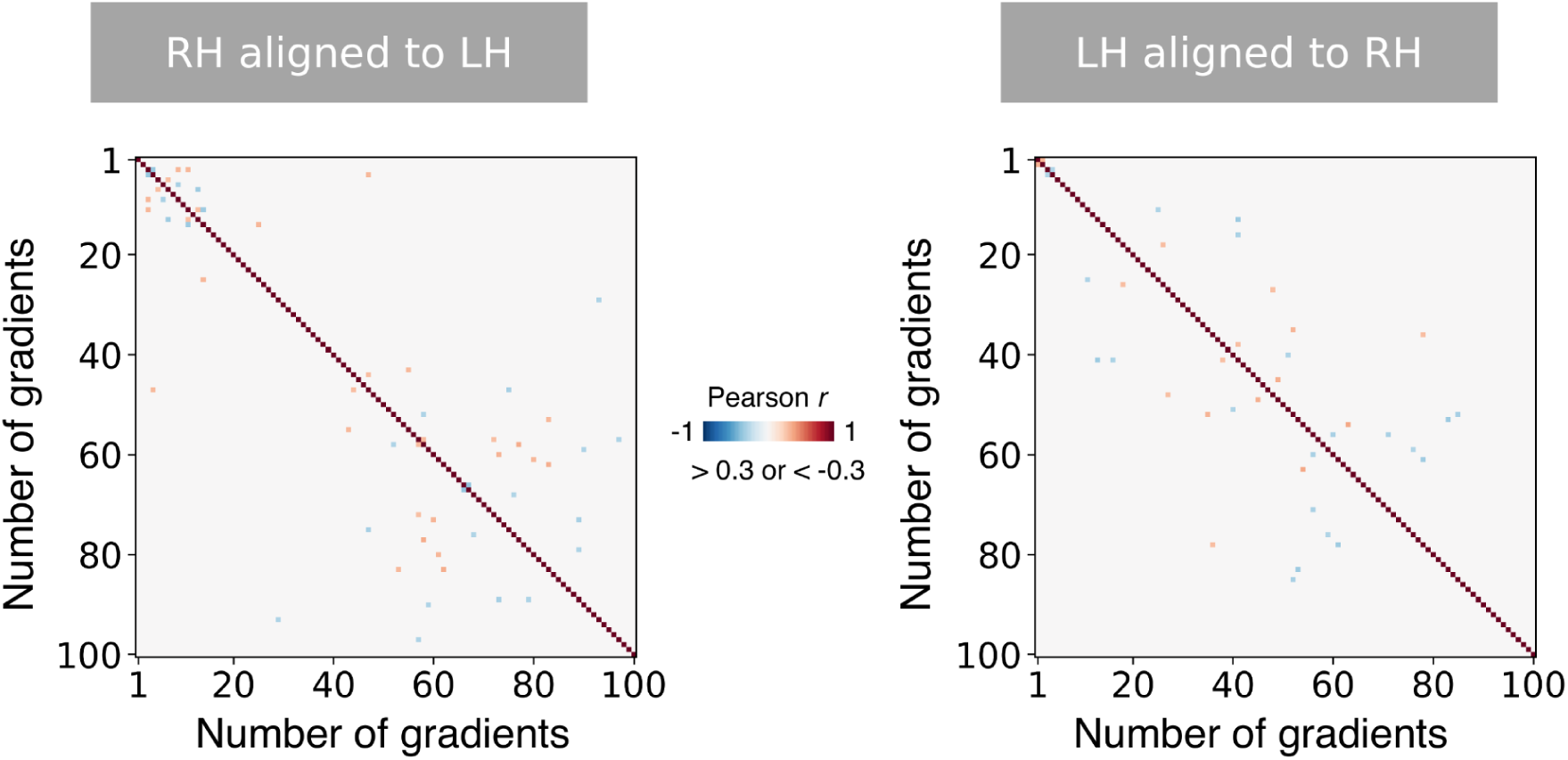
Interrelation between gradient asymmetry maps for different alignment template references. Left and right panels are LH and RH as the templates, respectively.

**Figure S7.**
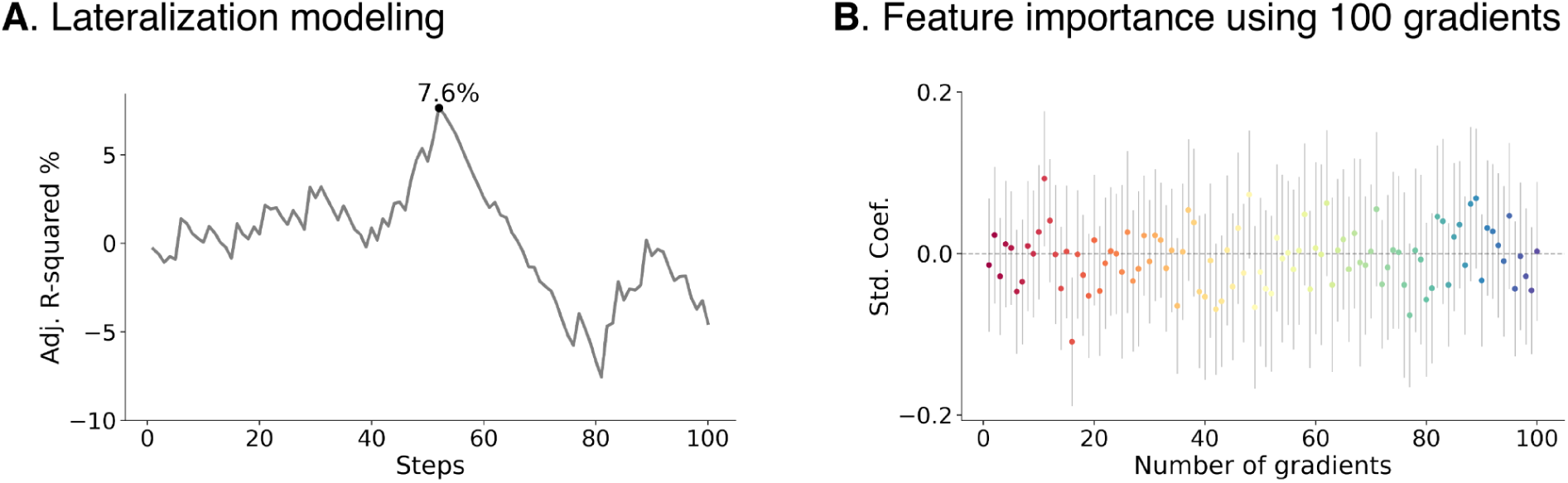
Asymmetry modeling when RH is the alignment template. **A.** Fitting curve along each step. **B**. Standardized regression coefficient and 95% confidence interval for asymmetry modeling.

**Figure S8.**
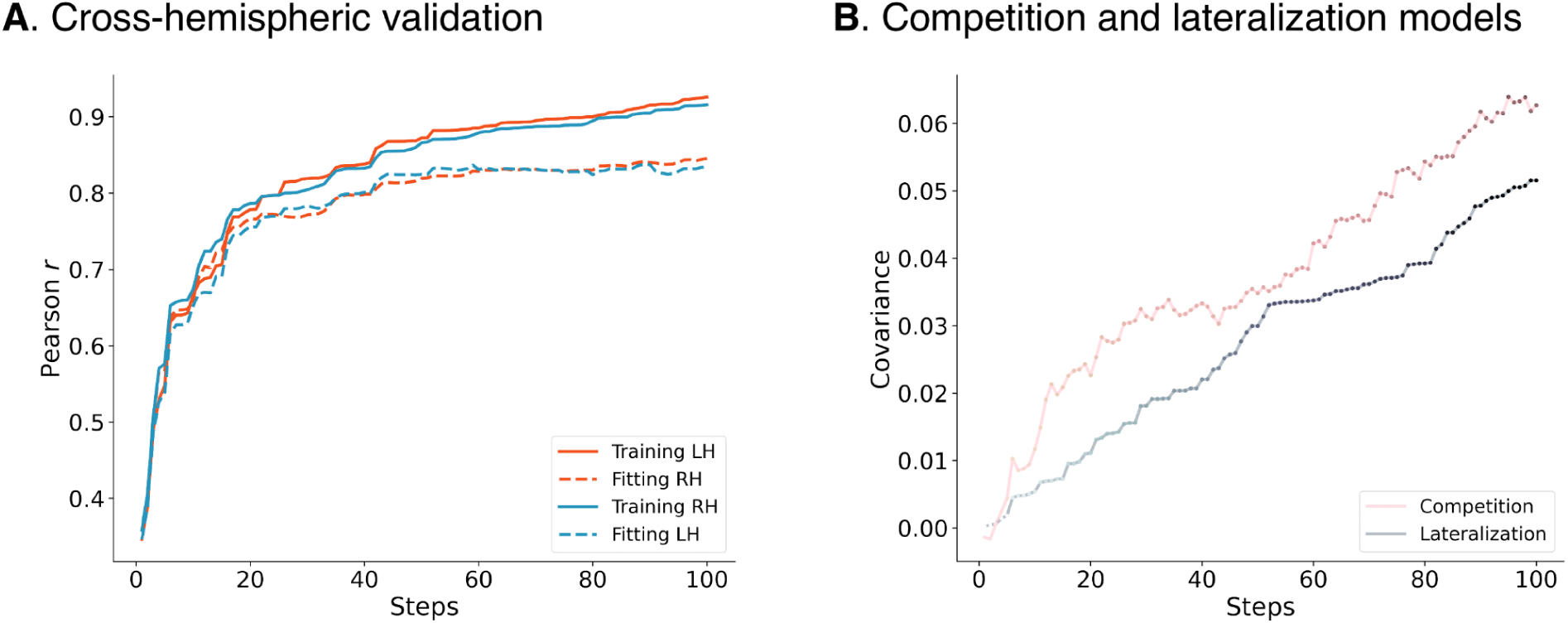
Cross-hemispheric validation when RH is the alignment template. **A.** Fitting curve for each hemisphere along each step. **B**. Competition versus lateralization models.

**Figure S9.**
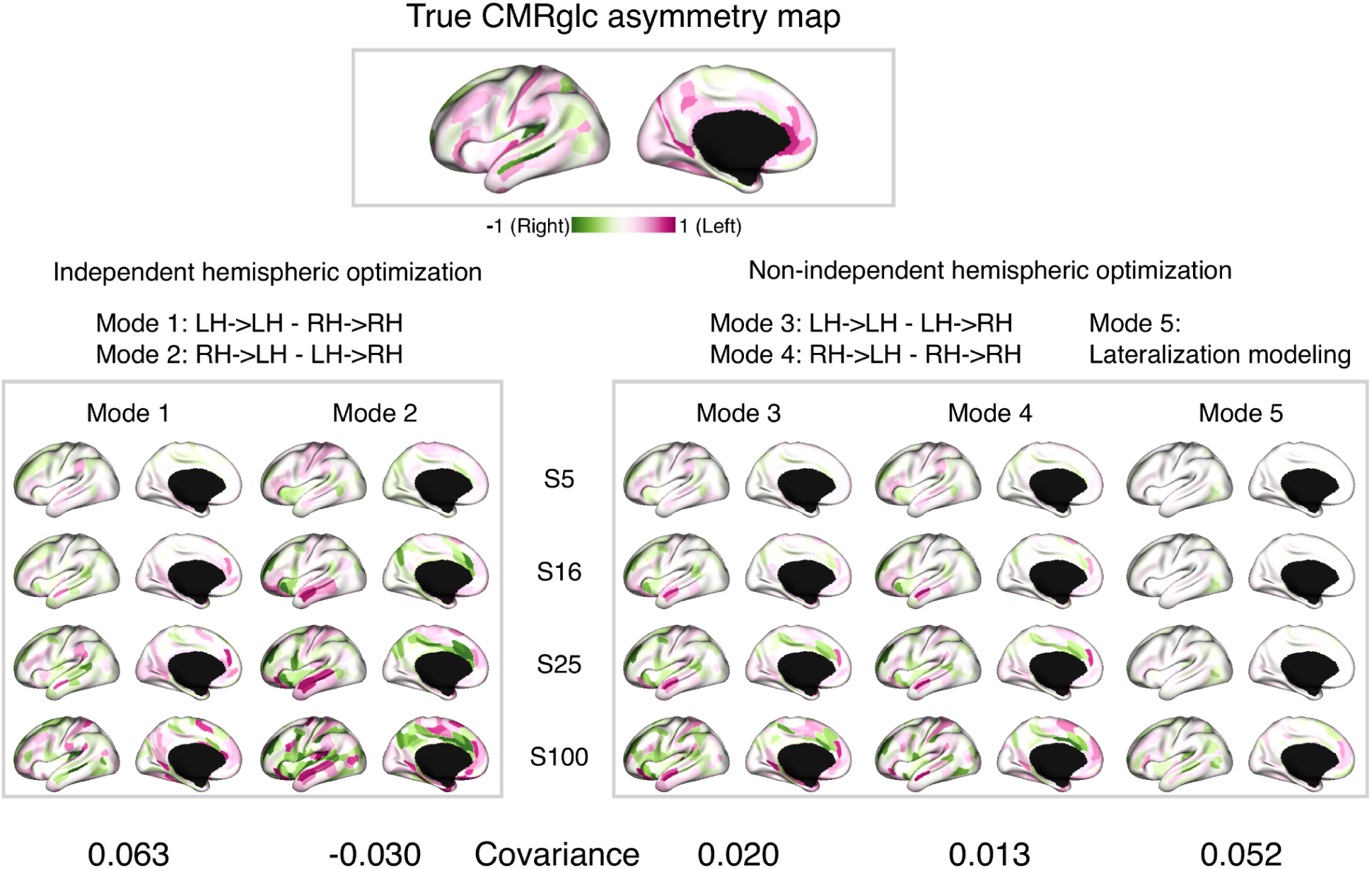
Competition versus lateralization using different modes.

**Figure S10.**
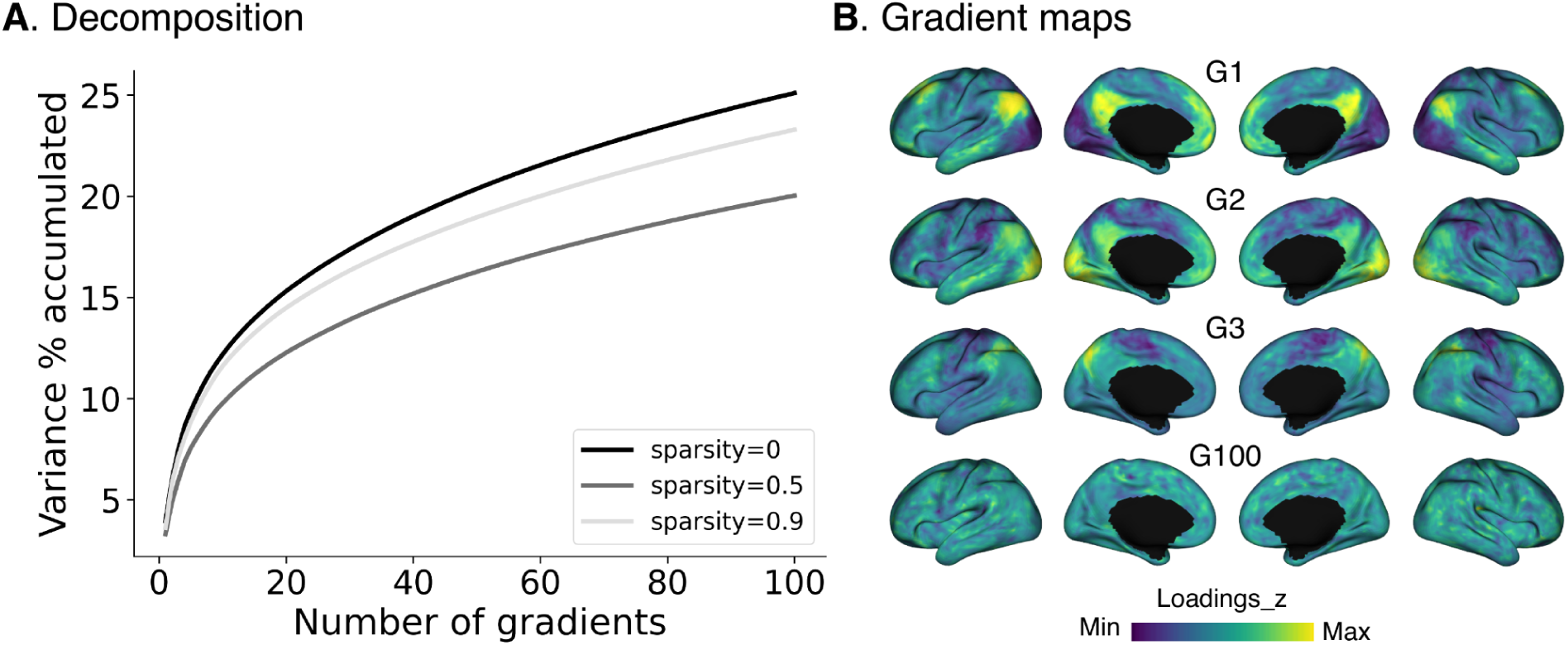
Analyses using fsLR-5k scheme (sparsity = 0, 0.5, and 0.9, separately). **A**. the accumulative lambda values for the decomposition. **B**. Gradient loadings of G1-3 and G100.

**Figure S11.**
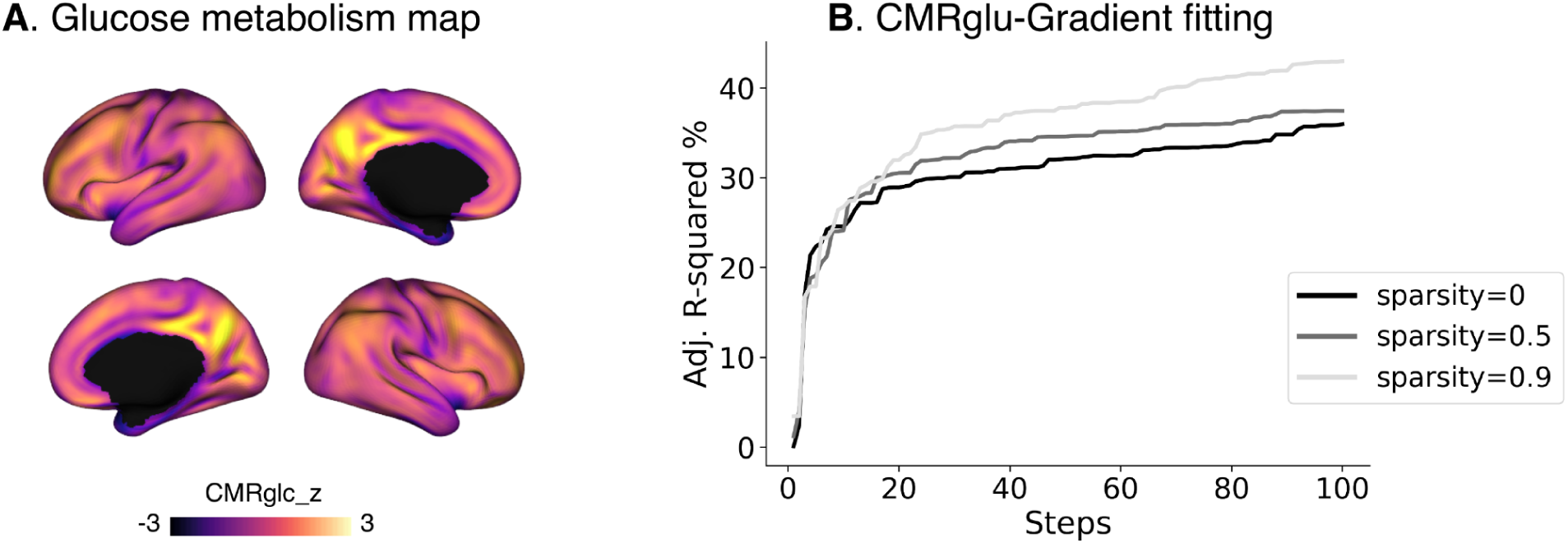
Glucose modeling using fsLR-5k scheme. **A**. Glucose metabolism (CMRglc) map. **B**. Fitting curve for sparsity = 0, 0.5, and 0.9, separately.

**Figure S12.**
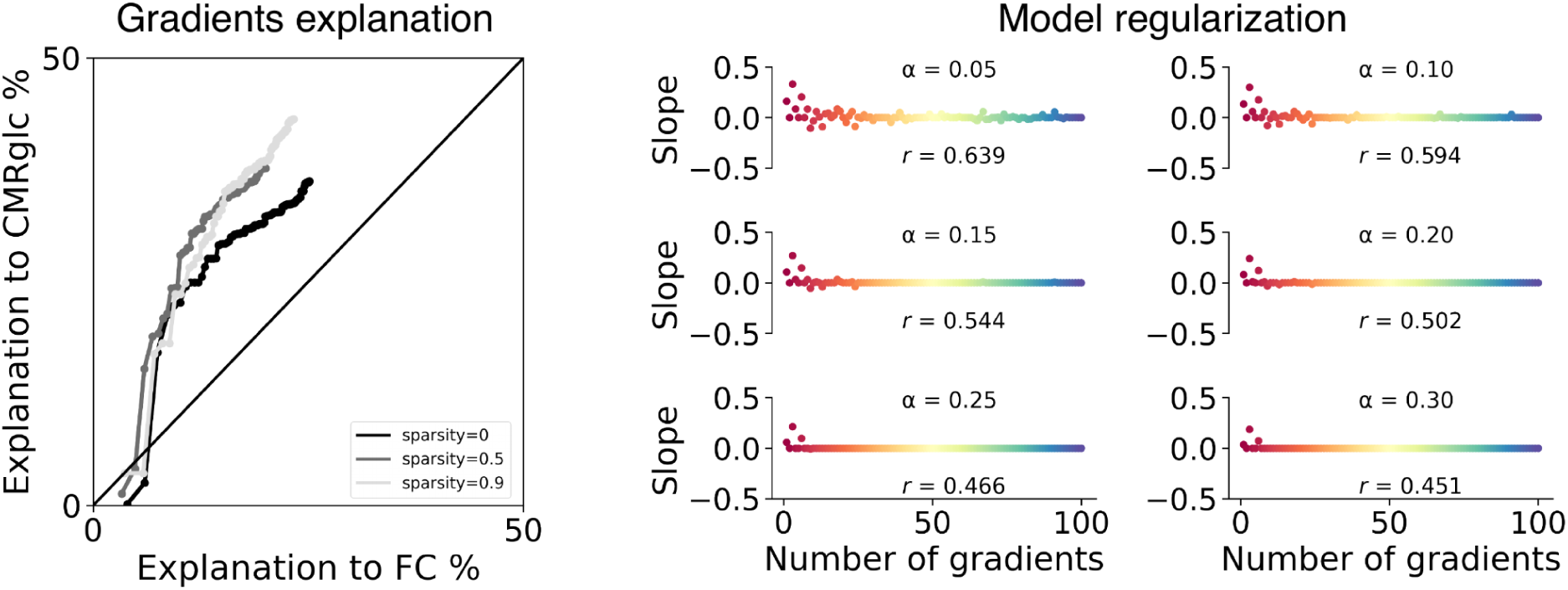
Gradient explanation to FC and CMRglc map and model regularization using fsLR-5k scheme.

**Figure S13.**
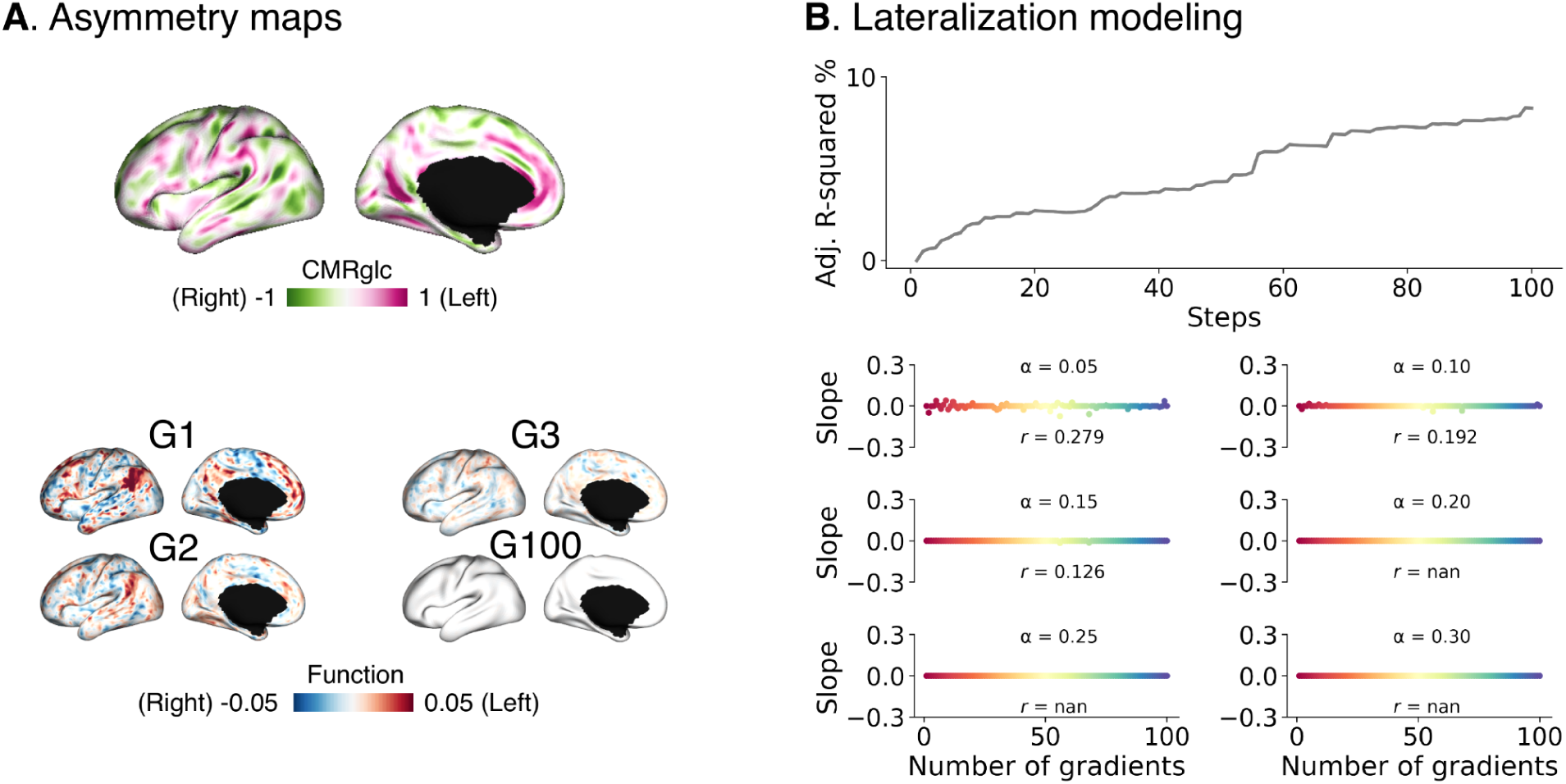
Asymmetry modeling using fsLR-5k scheme. **A**. Gradient asymmetry maps (G1-3 and G100). **B**. Fitting curve along steps and model regularization for step 100.

**Figure S14.**
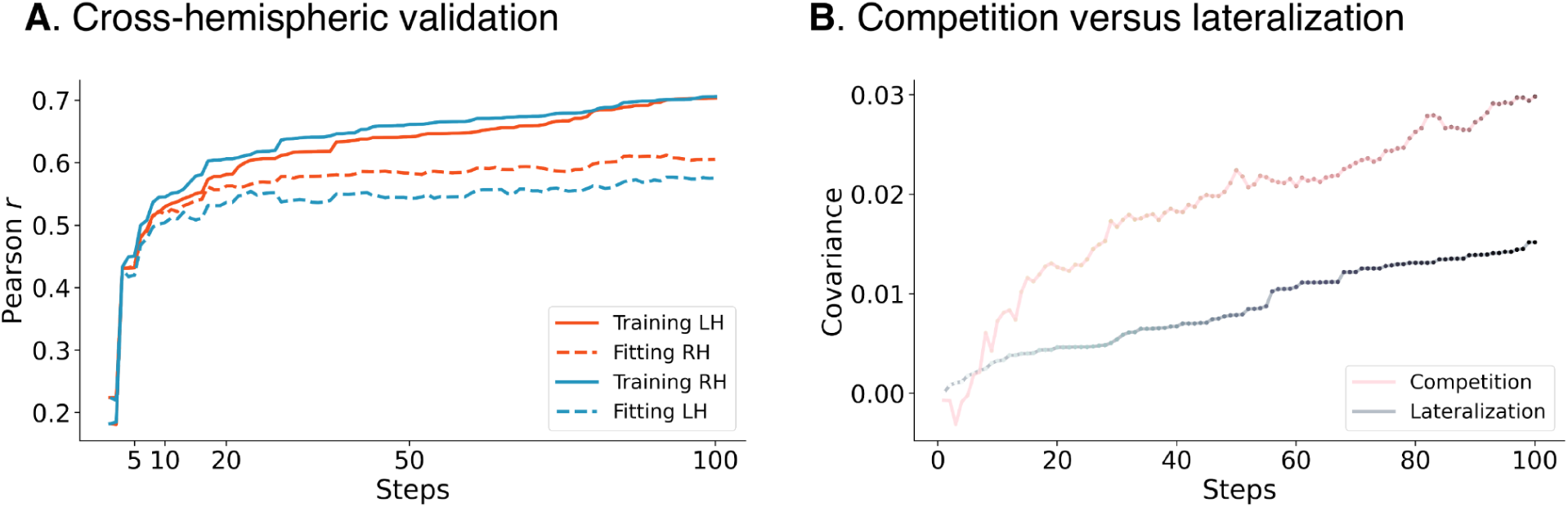
Cross-hemispheric validation using fsLR-5k scheme. **A.** Fitting curve for each hemisphere along each step. **B**. Competition versus lateralization models.

**Figure S15.**
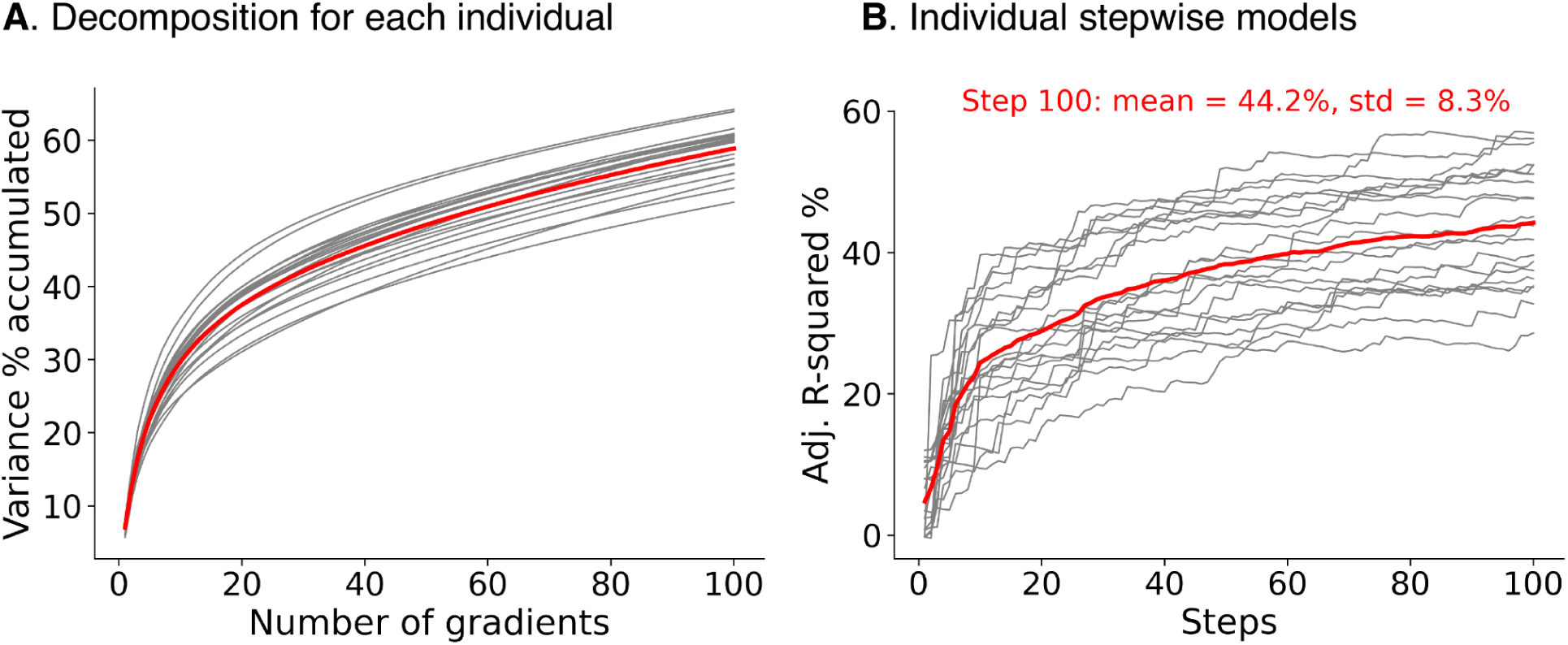
Individual analyses (sparsity = 0.9). **A**. the accumulative lambda values for the decomposition. **B**. Individual stepwise models. Gray and red lines indicate individual level curve and mean across individuals.

**Figure S16.**
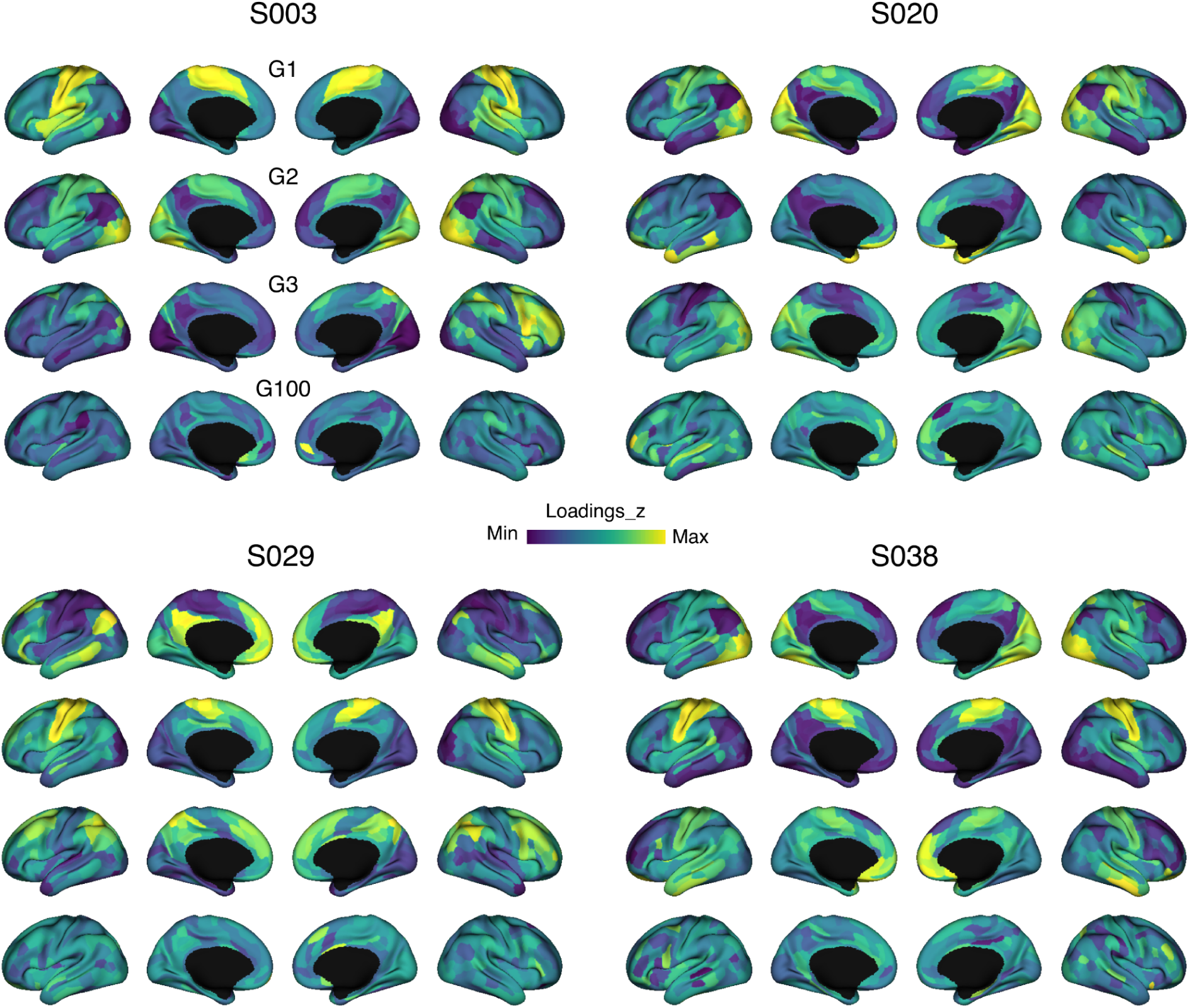
Raw gradient loading maps (G1-3 and G100) for four subjects (S003, S020, S029, and S038) .

**Figure S17.**
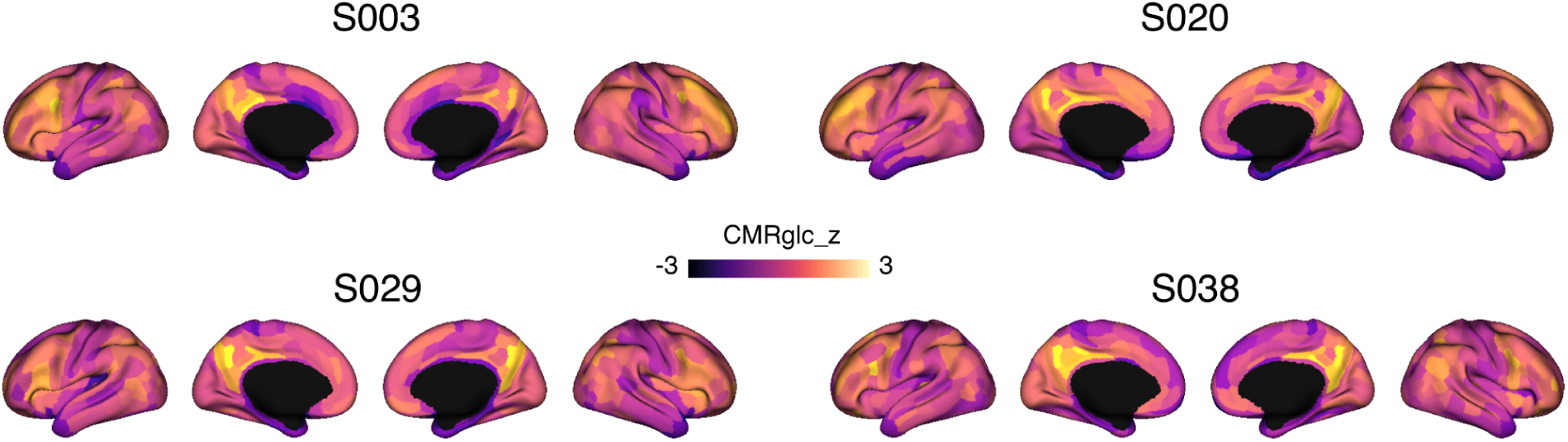
Raw glucose metabolism map (CMRglc) for four subjects(S003, S020, S029, and S038) .

**Figure S18.**
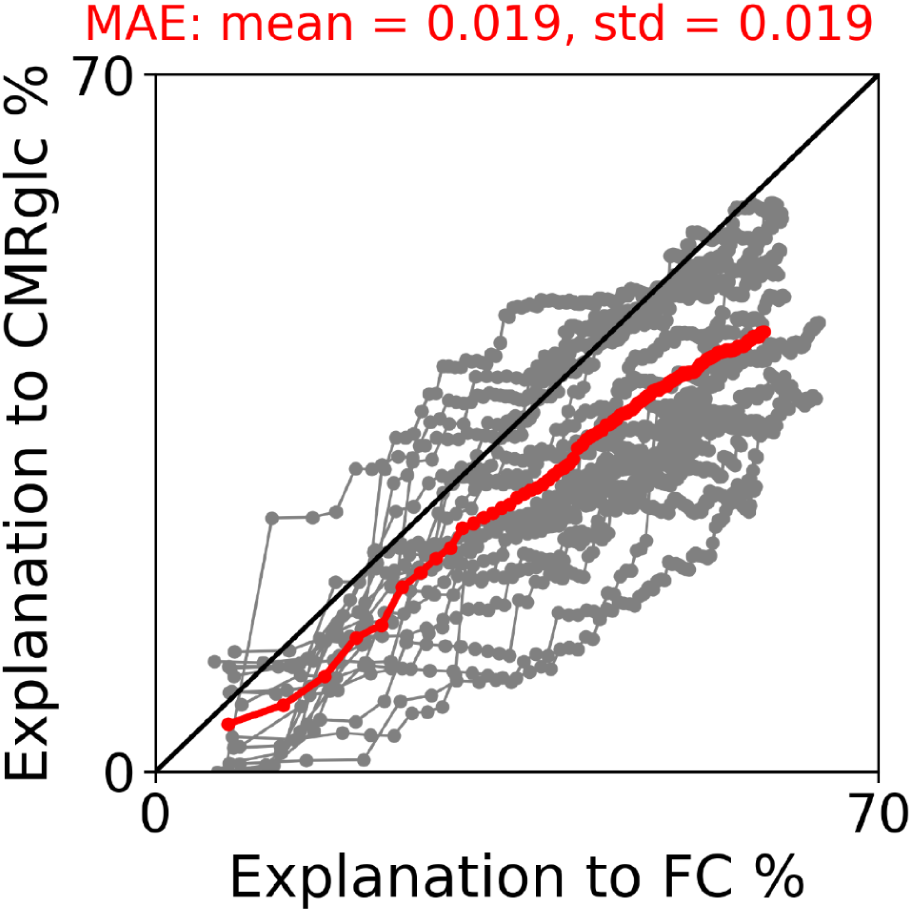
Gradient explanation to FC and CMRglu map at individual level. Gray and red lines indicate individual level curve and mean across individuals.

**Figure S19.**
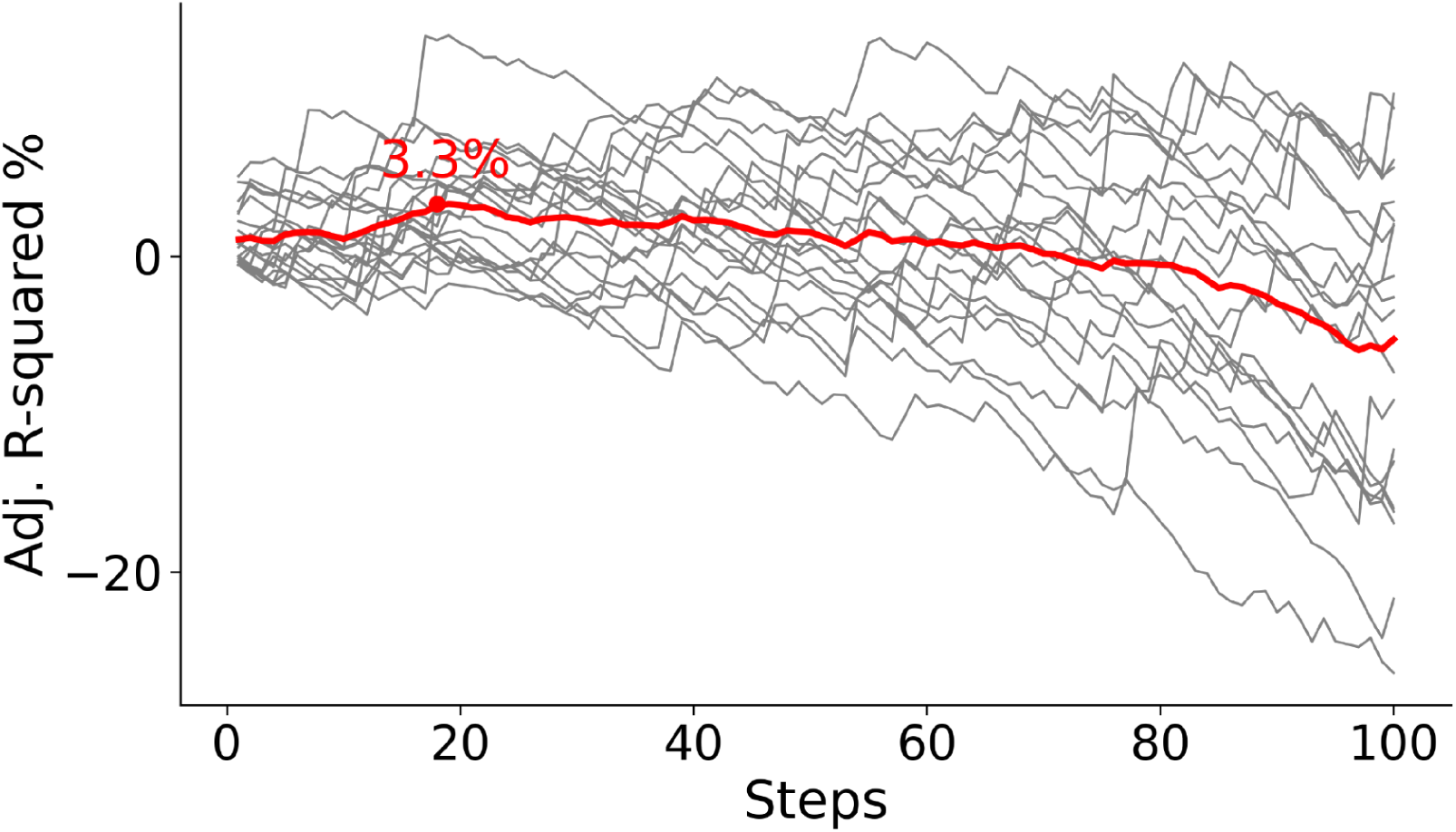
Asymmetry modeling at individual level. Gray and red lines indicate individual level curve and mean across individuals.

**Figure S20.**
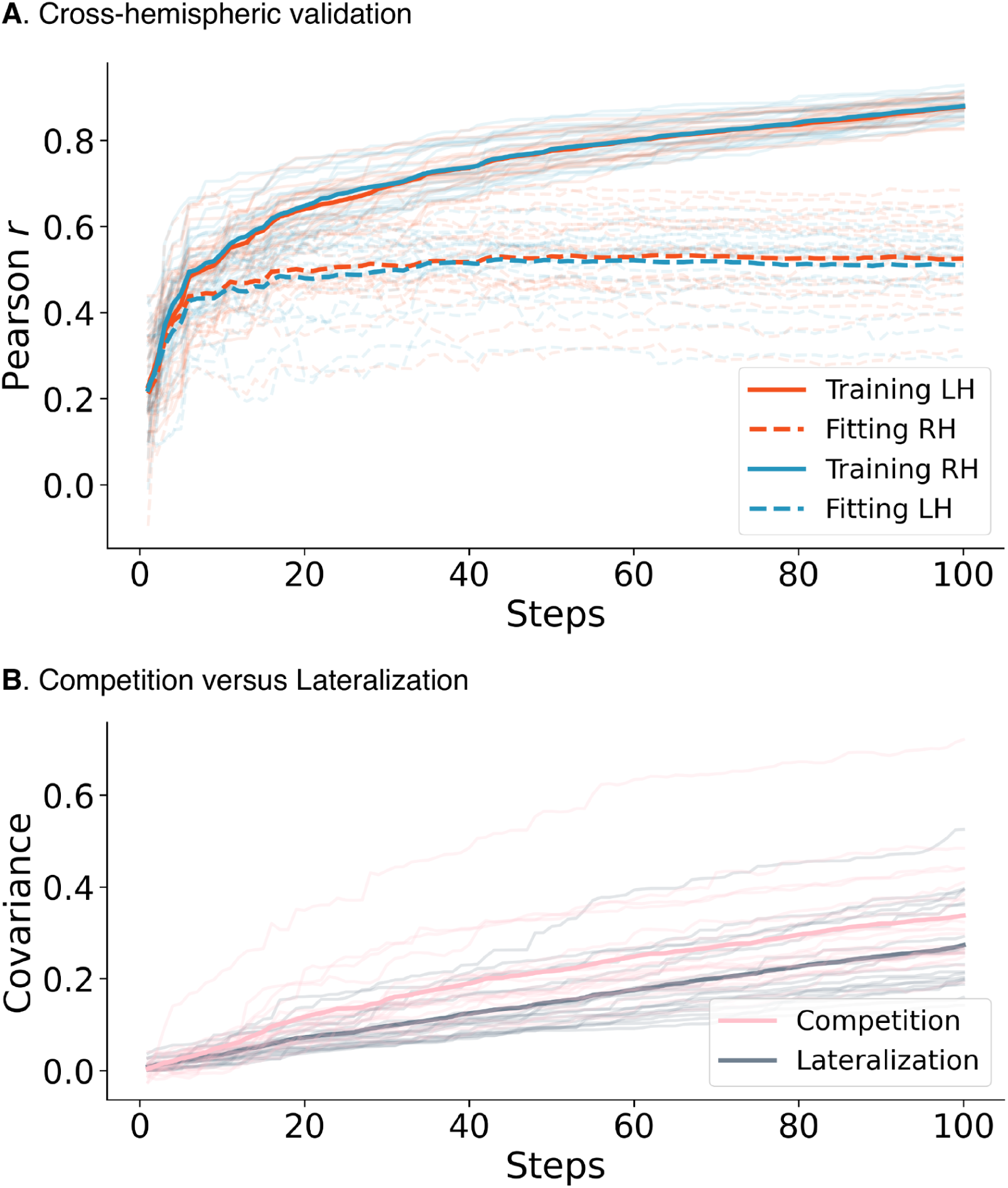
Cross-hemispheric validation (**A**) and competition versus lateralization (**B**) at individual level. Curves with transparency = 0.7 and 0 indicate an individual and mean across individuals.

**Figure S21.**
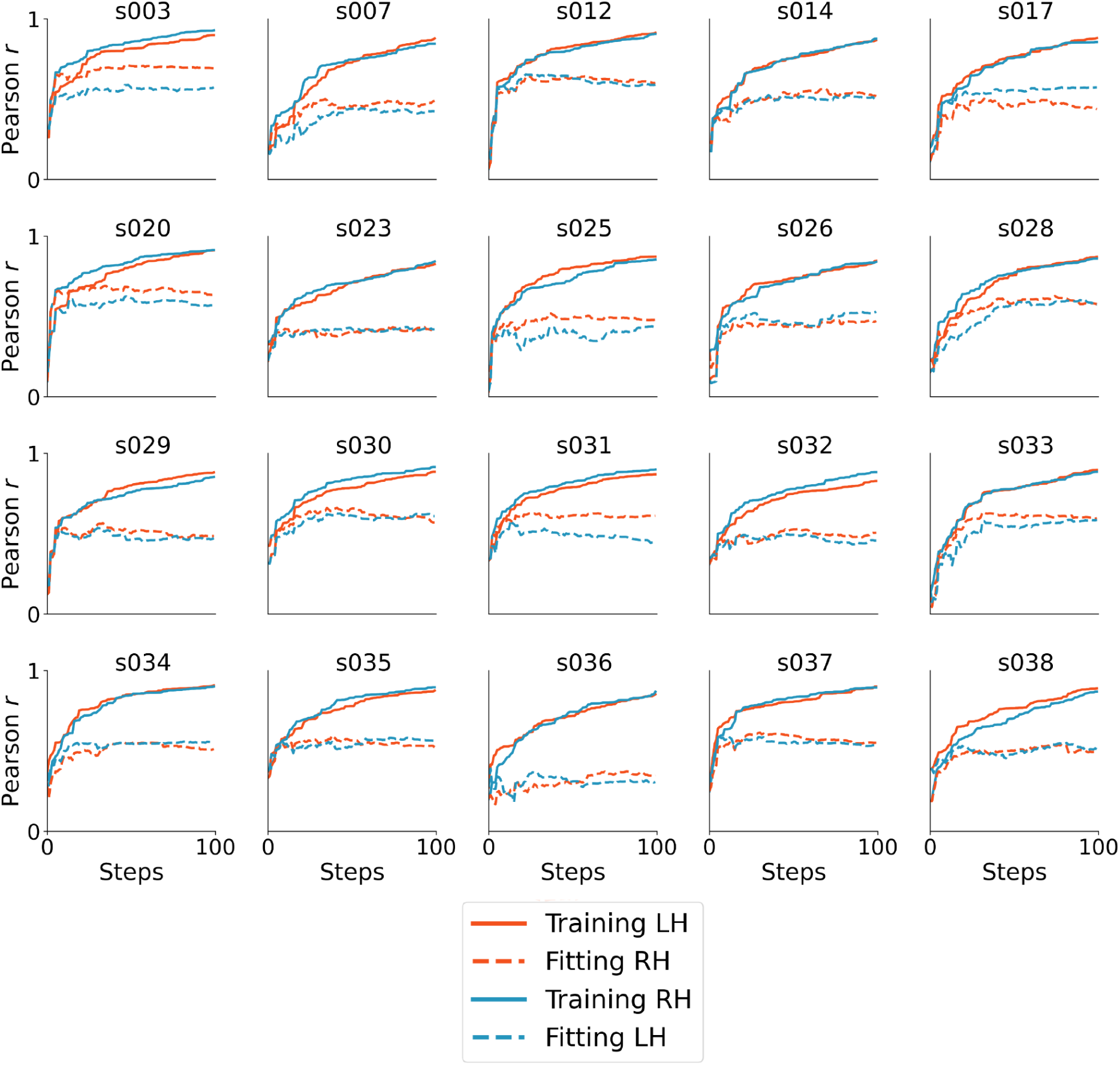
Cross-hemispheric validation for each individual.

**Figure S22.**
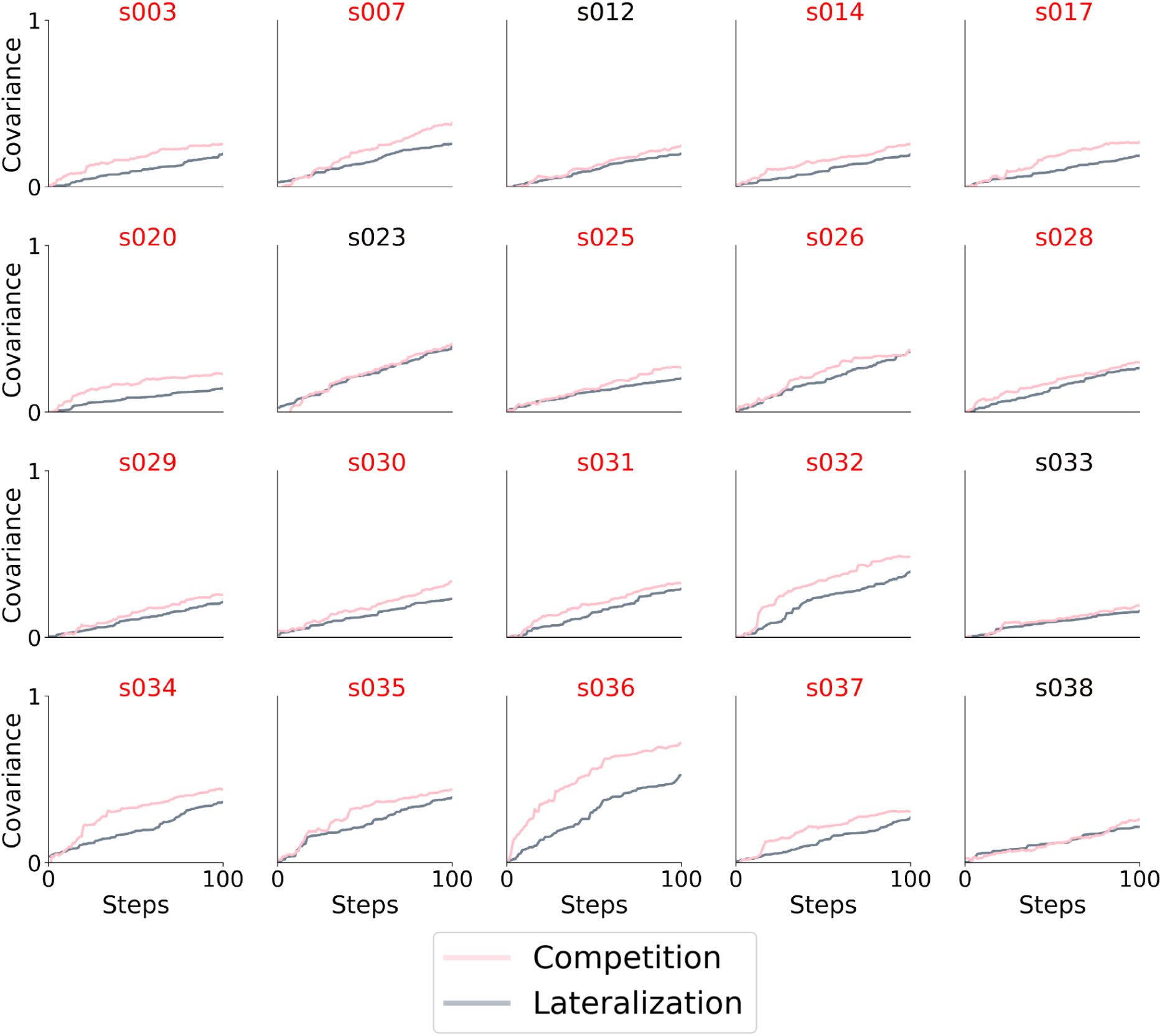
Competition versus lateralization for each individual. Red and black titles indicate individuals with clear and unclear comparison.

